# SMAD4 loss drives chromosomal instability during tumourigenesis via translational reprogramming

**DOI:** 10.64898/2026.03.23.712291

**Authors:** Julia V Milne, KaMeng Wu, Eric P Kusnadi, Sandra Brosda, Kenji M Fujihara, Ebtihal H Mustafa, Sasha Witts, Katherine Papastratos, Maree Pechlivanis, Anna S Trigos, Metta K Jana, Paul J McMillan, Thomas D Jackson, Niko Thio, Yuzhou Feng, Karen G Montgomery, John Alexander, Mark Wade, Andrew P Barbour, Kaylene J Simpson, Cuong P Duong, David S Liu, Wayne A Phillips, Luc Furic, Nicholas J Clemons

## Abstract

Chromosomal instability (CIN), arising from errors in chromosome segregation during cell division, is a hallmark of cancer. Whilst CIN can result from several mitotic defects, the mechanisms that initiate CIN to drive tumourigenesis remain incompletely understood. Here, we show that loss of SMAD4 reprograms translation to induce CIN, resulting in tumour formation. Multi-omics analysis of tumourigenesis models driven by loss of SMAD4 complemented by functional studies demonstrate that loss of SMAD4 in pre-neoplastic cells leads to dysfunctional mitosis and an altered global translation landscape. We show that CDK11B is translationally downregulated in *SMAD4^-\-^* cells, and re-expression of the mitosis-specific isoform of this protein (CDK11B-p58) rescues the mitotic defects. Analysis of patient tumours reveals a strong correlation between markers of CIN and SMAD4 status, indicating the clinical relevance of this phenotype. Collectively, we reveal a previously unrecognised role for SMAD4 as a gatekeeper for CIN-mediated tumourigenesis via regulation of translation.

## Introduction

Over a decade ago, genome instability was designated a tumour-enabling hallmark of cancer (1). Unstable genomes present with a wide range of characteristics, including chromosome segment alterations or fusions, ploidy abnormalities, whole genome duplications, and gene mutability, which frequently drive tumour evolution and heterogeneity. Chromosomal instability (CIN) is the predominant form of genome instability, and may arise from a multitude of errors during cell division, such as chromosome mis-segregation, defects in the mitotic spindle assembly checkpoint, or centrosome aberrations (2, 3). Many factors, including cyclin-dependent kinases (CDKs), play integral roles in the progression of proper mitoses (3, 4). Thus, impairment of any of the numerous elements involved in these complex processes may result in CIN.

Most cancers exhibit some extent of CIN (5, 6), and there are many models of CIN driving tumourigenesis (7–9) and resistance to therapy (10, 11). In particular, cancers of the gastrointestinal (GI) tract are characterised by CIN phenotypes (12–14). For example, oesophageal adenocarcinoma (OAC) develops from the precursor metaplasia, Barrett’s oesophagus (15); a progression that correlates with increasing CIN (16, 17). We have previously shown that SMAD4 loss, which coincides with the transition from metaplasia to OAC, induces tumourigenesis and increased copy number alterations (CNA) in Barrett’s oesophagus cells (18). The observed CNA increase was especially pronounced in the resultant tumour cells, suggesting that accumulation of mitotic errors occurs in these cells throughout their oncogenic progression.

Loss of SMAD4 is common in many solid tumour types, particularly in pancreatic and other GI cancers (19, 20). Consequences of SMAD4 loss are frequently attributed to loss of canonical TGFβ and BMP signalling, producing effects such as suppression of growth arrest pathways and altered regulation of epithelial-to-mesenchymal transition (EMT) (21, 22). One recent study showed that SMAD4-deficient colorectal cancer cells exhibit irregular mitotic spindle formation (23), a phenotype that had not previously been associated with SMAD4 loss but has not been further explored. Combined with our previous findings, this led us to hypothesise that SMAD4 loss may lead to CIN through deregulation of mitosis.

In this study, using unbiased approaches we demonstrate that SMAD4 plays a key role in maintaining chromosomal integrity and suppressing tumourigenesis. We discovered that SMAD4-deficient cells deregulate translation, resulting in altered mitosis and accumulation of CIN. These findings point to a previously undiscovered role for SMAD4 in maintaining genomic stability to suppress tumourigenesis, that is distinct from its canonical functions.

## Results

### SMAD4 loss is associated with chromosomal instability and a deregulated mitosis signature

We first aimed to understand the broad genomic consequences of SMAD4 deficiency in OAC and other GI cancers with frequent *SMAD4* mutation. To do this, we interrogated patient datasets from The Cancer Genome Atlas (TCGA) Firehose Legacy and Memorial Sloan Kettering (MSK) studies (13, 24–26). Given its prevalence in OAC and known role in CIN (27, 28), we stratified samples by presence of *TP53* alteration (i.e., any mutation or CNA) or *SMAD4* alteration and assessed the fraction of genome alteration (FGA). We identified a significant change in FGA only as a function of *SMAD4* alteration in *TP53* altered cells, with a trend towards the same in *TP53* unaltered samples (Figure 1A). Unexpectedly, alteration of *TP53* was not associated with an increase in FGA compared to *TP53* wildtype, unless in combination with *SMAD4* alteration. Similar patterns were detected in other GI cancers, including gastric, colorectal and pancreatic adenocarcinoma (Figure S1A-C). Interestingly, three out of five of the OAC patient samples with wildtype p53 and altered *SMAD4* were also found to have *MDM2* copy number gains, whilst a further sample expressed low levels of p53 protein, comparable to that of *MDM2*-amplified samples (Supplementary Table S1). Additionally, significant associations were identified between *TGFBR2* or *MYC* alterations and the fraction of the genome altered (Figure S1D), indicating that this phenotype may potentially be extended to loss of other TGFβ pathway genes or downstream TGFβ targets. Combined, these findings suggest that SMAD4 status is a stronger determinant of proportion of genomic alteration than that of *TP53* in OAC, and a strong determinant in other GI cancer types.

**Figure 1.**
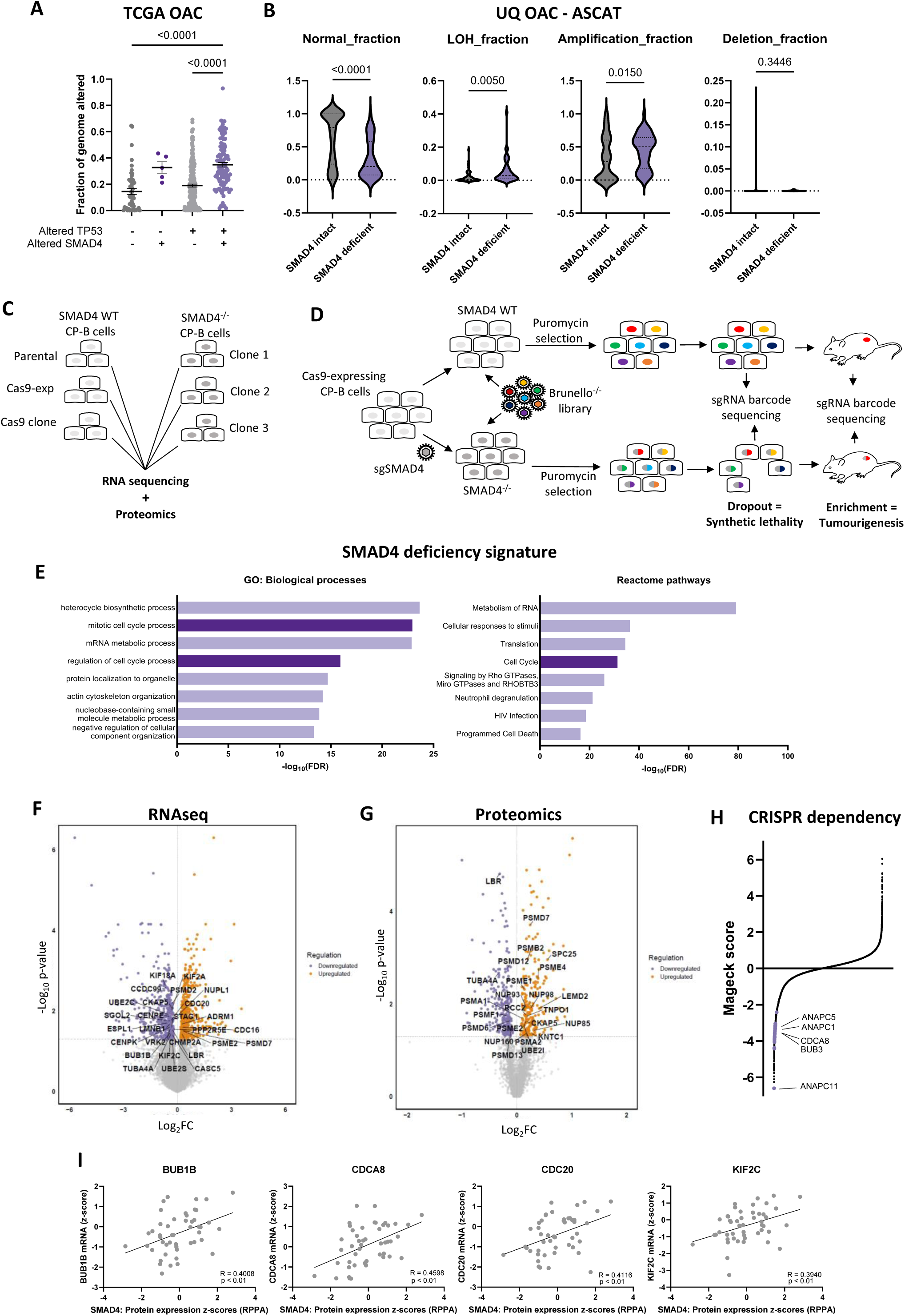
SMAD4 loss is associated with chromosomal instability and a deregulated mitosis signature in OAC. (A) Dot plots showing fraction of the genome altered in TCGA and MSK OAC patient samples as a function of alteration of *SMAD4* and *TP53* genes. Error bars represent SEM. (B) Violin plots indicating median and quartiles of chromosomal aberrations in UQ OAC patient samples, stratified by *SMAD4* status. Schematics showing the workflow for the (C) RNA-sequencing and proteomics experiments and (D) *in vitro* CRISPR-Cas9 knockout screen using Brunello genome-wide knockout library (Brunello^-/-^). (E) Bar charts depicting top significant Biological processes and Reactome pathways in SMAD4 deficient cells. Volcano plots depicting differentially expressed (F) genes and (G) proteins between CP-B parental and CP-B *SMAD4*^-/-^ cells as determined by RNA sequencing and LC-MS/MS, respectively. (H) Scatter plot depicting ranked MaGeCK scores, where <0 = dropout and >0 = enrichment of knockout. (I) Scatter plots depicting mRNA expression z-score of genes as a function of SMAD4 protein expression z-score in TCGA and MSK OAC patient samples. (A) Ordinary one-way ANOVA with Tukey’s multiple comparisons test, (B) unpaired Student’s t-test. (I) Correlation determined by simple linear regression and Pearson’s R.

To more deeply evaluate the relationship between SMAD4 status and genomic alterations (specifically, chromosomal abnormalities) we examined an independent set of OAC patient samples from which we obtained WGS, WES and/or SNP array with matched RNAseq data (University of Queensland dataset; UQ) (29, 30). Samples were designated SMAD4-deficient or SMAD4-intact based on copy number, mutation and gene expression status (Table S2). From a cohort of 295 patients, 54 samples were considered “deficient” and 116 “intact”. Where available, relative mRNA expression data from RNAseq were used to verify SMAD4 expression levels in cases of uncertainty (Figure S1E), and ambiguous samples were excluded from analysis. We investigated these sample sets (“intact” versus “deficient”) for patterns of genomic alterations that may be influenced by SMAD4 status using allele-specific copy number analysis of tumours (ASCAT) (31). SMAD4-deficient samples exhibited significantly more CNA than SMAD4-intact samples (as indicated by a lower normal fraction) and were significantly more likely to exhibit LOH and amplifications, without complete gene deletion (Figure 1B). This suggests a tendency for allelic loss or gain via single chromosome alteration. Because classification of SMAD4 status could vary depending on the chosen definition of deleterious alteration, we performed complementary analyses using more strict criteria (“strict deficient” and “strict intact”), which corroborated our findings (Figure S1F). We also assessed samples using a homologous recombination deficiency (HRD) score (32) as an indication of genomic instability. SMAD4-deficient samples displayed increased telomeric allelic imbalance (TAI), LOH, large scale transitions (LST) and a higher overall HRD score (Figure S1G). To eradicate the possibility that chromosomal loss of SMAD4 itself may be confounding the data, we repeated these analyses after removing chromosome 18q21.2 and found no change in significant features (Figure S1H). Strikingly, we found that all but one of the “strict deficient” samples had concurrent alterations in *TP53* (Figure S1I). Amplification of MDM2 was detected in the one sample with wildtype TP53 (Table S1), supporting the idea that SMAD4 deficiency may only occur on a background of TP53 loss or mutation. These data confirm that the observed CIN phenotype is dependent on *SMAD4* status and not due to segregation of *TP53* mutation between the two groups.

We next employed complementary ‘omics approaches to obtain a global view of the effect of SMAD4 loss on the transcriptome, proteome and gene dependency of a non-tumourigenic Barrett’s oesophagus cell line (CP-B) (33), which we had previously shown becomes tumourigenic upon SMAD4 loss (18). We obtained significant differentially expressed gene and protein sets from 3’ RNA-sequencing (RNAseq) and label-free quantitative proteomics, respectively, of parental and CRISPR-Cas9-mediated *SMAD4*^-/-^ CP-B cells (Figure 1C). We also conducted a genome-wide CRISPR-Cas9 screen to identify gene knockouts that were selectively lethal in the *SMAD4*^-/-^ cells (i.e., *synthetic lethal* hits) (Figure 1D). The gene lists from RNAseq, proteomics and the CRISPR dependency screen were collectively analysed using the Metascape resource (34) to identify overlapping genes and gene ontologies (GO). This analysis revealed multiple significantly enriched GO clusters in SMAD4-deficient cells, including processes and pathways involved in mitosis (Figure 1E, Figure S2A-B). Of particular interest were enriched pathways pertaining to chromosome segregation and the transition of mitotic metaphase to anaphase, strongly suggesting a deregulation of these processes in SMAD4-deficient cells. Importantly, a number of genes within these processes (namely *TTK*/MPS1, *UBE2S*, *CENPE*, *SPDL1*, *BUB1B*, *UBE2C*, *CDC16*/ANAPC6, *CDC20* and *CDCA8*) are implicated in the spindle assembly checkpoint (SAC) and anaphase-promoting complex/cyclosome (APC/C) (35), and were significantly downregulated in *SMAD4*^-/-^ cells (Figure 1F). At the protein level, *SMAD4*^-/-^ cells showed deregulation in this same metaphase-to-anaphase transitional process, with particular perturbation to proteasome subunits required for correct APC/C functionality (Figure 1G) (36, 37). Meanwhile, the genome-wide knockout screen revealed that SMAD4-deficient cells were dependent on several genes involved in the APC/C (*ANAPC11*, *ANAPC1*, *ANAPC5*) and the SAC (*BUB3*, *CDCA8*) and on the cell cycle checkpoint kinase, CHEK1 (Figure 1H). We confirmed increased dependency of *SMAD4^-/-^* cells on ANAPC11 and CHEK1 using genetic or pharmacological approaches, respectively (Figure S2C-D). These data lend weight to the notion that SMAD4 loss deregulates mitotic processes (in particular, the metaphase-to-anaphase transition) and engenders an increased dependency on specific components of these processes for cell proliferation or survival.

In the TCGA OAC patient samples, we found significant positive correlations between SMAD4 protein levels and mRNA expression of *BUB1B*, *CDCA8*, *CDC20* and *KIF2C* (Figure 1I), despite no previously known relationship between SMAD4 and these genes. Similar findings were identified upon examination of the Cancer Cell Line Encyclopaedia (CCLE) mRNA expression and proteomics data on the Cancer Dependency Map (DepMap) portal (38). This analysis revealed positive correlations between SMAD4 protein levels and the mRNA and protein levels of *BUB1B*, *CDCA8*, *SGO1*, *NDC80* and *FBXO8* in pan-cancer cell lines (Figure S2E-F). These data point towards a broader implication of SMAD4 loss on expression of mitotic spindle genes and proteins.

Specific gene aberrations have been associated with increased chromosomal instability (39) and mitotic abnormalities (40), and one recent study identified a dysregulated mitotic spindle gene expression signature associated with oesophageal cancer (41). We next examined the overlap of these gene sets with our differentially expressed genes following SMAD4 knockout. Again, utilising Metascape analysis, we identified many genes and gene ontologies that co-occurred in our dataset and the previously published CIN and mitotic spindle signature gene sets (Figure S2G). The most significantly enriched cluster of gene ontologies pertained to “mitotic metaphase and anaphase”, signalling a common deregulation of these mitotic phases amongst the datasets. Scrutinising these datasets individually, we identified statistically significant overlap between our SMAD4-deficiency signature and the CIN70 signature (39) (representation factor: 3.4; p < 1.83 x 10^-4^) as well as the oesophageal cancer-specific dysregulated mitotic spindle signature (41) (representation factor: 4.6; p < 3.65 x 10^-9^). Taken together, these data suggest that our SMAD4-deficiency signature may indeed reflect a deregulated mitosis signature.

### Loss of SMAD4 increases mitosis defects in pre-neoplastic Barrett’s oesophagus cells

To investigate this potential deregulated mitosis phenotype, we performed live-cell confocal, time-lapse imaging of CP-B parental and *SMAD4*^-/-^ cells, as well as an additional Barrett’s oesophagus cell line, CP-D, from which we derived further *SMAD4*^-/-^ populations (Figure S3A). Chromosome dynamics were visualised using a DNA stain and images recorded at eight-minute intervals for 16-18 hours (Figure 2A-C). As expected, due to endogenous *TP53* mutations in these cell lines, a fraction of each of the parental cell lines exhibited defects in mitosis such as anaphase bridges, lagging chromosomes, failed cytokinesis or mitotic slippage, whilst many progressed through mitosis without error. Interestingly, the percentage of mitotic defects observed in *SMAD4*^-/-^ CP-B lines was greater than respective parental cells, but no overall difference in total defects was observed in CP-D cells with or without SMAD4 (Figure 2D-E). However, *SMAD4*^-/-^ in both cell lines led to an increased proportion of mitoses with lagging chromosomes, resulting in micronuclei in the daughter cells, and a trend towards more multinuclear cells (Figure 2F-G). Interestingly, there was no difference in mitotic index (Figure S3B), nor in time spent in mitosis overall or from nuclear envelope breakdown to anaphase (Figure S3C-D). This suggests similar rates of cell division and no induction of mitotic checkpoints that might prolong mitotic duration in SMAD4-deficient cells.

**Figure 2.**
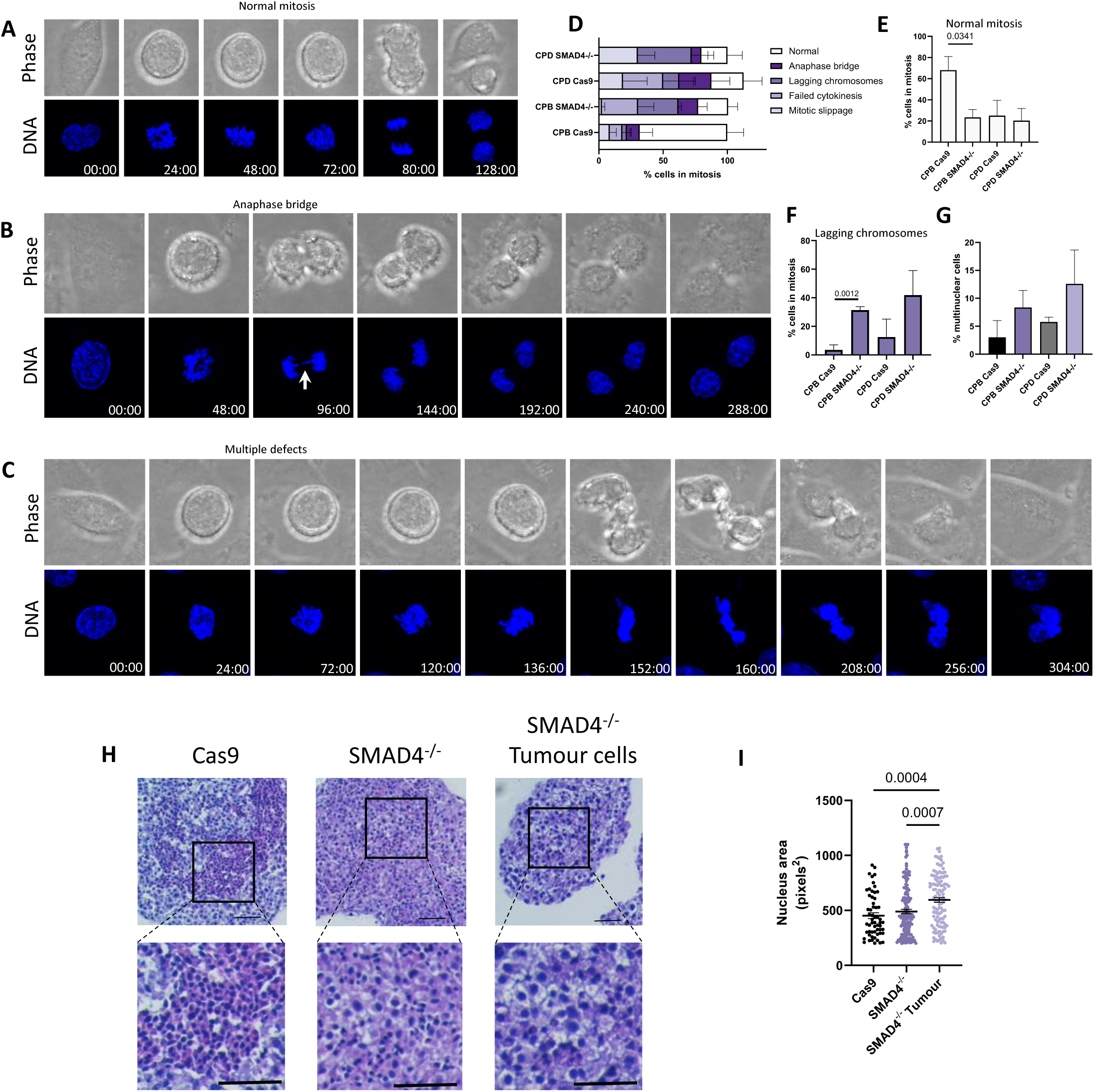
Loss of SMAD4 increases mitosis defects in Barrett’s oesophagus cells. Time lapse images of CP-B *SMAD4*^-/-^ cells stained with SPY650-DNA showing examples of (A) normal mitosis, (B) lagging chromosomes and (C) failed cytokinesis. Bar charts showing percentage of (D) mitotic cells with mitosis defects (range 9-38 cells in mitosis visualised per cell line), percentage of cells with (E) normal mitoses, (F) lagging chromosomes, and (G) multiple nuclei (87-170 cells visualised per cell line). (H) Haematoxylin and eosin staining of CP-B cells grown in 3-dimensional spheroids. Scale bars represent 50 µm. (I) Quantitation of nuclei area in CP-B cell spheroids. Solid bar represents mean, error bars represent SEM. Ordinary one-way ANOVA with Tukey’s multiple comparisons test.

We analysed the size of nuclei in CP-B parental and *SMAD4*^-/-^ cells, as well as a new cell line derived from *SMAD4*^-/-^ tumours that grew from xenografts of these cells (Figure S3A) (18). The cell lines were grown into spheroids, from which cross sections were analysed for nuclei size (Figure 2H-I). As expected, the size and variability of nuclei size increased across the cell line spectrum, where the parental CP-B cells displayed relatively small and uniform nuclei (mean = 452 pixels^2^, range = 202-912, SD = 197) compared to the *SMAD4*^-/-^ (mean = 490 pixels^2^, range = 200-1099, SD = 234) and tumour-derived cells, the latter of which exhibited very large nuclei (mean = 595 pixels^2^, range = 204-1065, SD = 234). This increase in size and variability of nuclei suggests a concordant increase in mitotic defects leading to progressive aberrant chromosome numbers (42) during cell transformation.

### SMAD4 loss and mTOR deregulation co-operate to drive tumourigenesis

Our tumourigenesis model of SMAD4 deficiency is characterised by a period of latency before tumour initiation, whereas there was no latency when cells derived from these tumours were injected subcutaneously into a subsequent generation of mice (18). We therefore theorised that the cumulative mitotic defects associated with SMAD4 loss may result in specific genomic alterations that would contribute to tumourigenic potential. Thus, we sought to identify potential tumour suppressor genes whose loss may co-operate with that of SMAD4 to accelerate tumour onset. We conducted a pooled genome-wide *in vivo* CRISPR-Cas9 (Brunello^-/-^ library (43)) enrichment screen in CP-B cells, on a background of either prior CRISPR-Cas9 mediated *SMAD4* knockout or SMAD4 wildtype (Figure 1D). This screen resulted in tumour initiation four-fold faster than our original model of *SMAD4*^-/-^-induced tumourigenesis, with palpable tumours noted after a mean of 21 days versus 160 days with *SMAD4*^-/-^alone (Figure 3A). In one case, Cas9-only expressing CP-B cells transduced with the genome-wide library also formed a tumour, however the time to tumour onset (273 days) was substantially longer than the *SMAD4*^-/-^ population. All tumours of sufficient volume were sequenced to identify the sgRNAs present in the population and their enrichment compared to the representation of sgRNAs in the cells before xenotransplantation. sgRNAs targeting *PTEN*, *TSC2* and *NF2*, notably all genes with downstream negative regulation effects on the PI3K/Akt/mTOR pathway (44–46), were all identified in the *SMAD4*^-/-^Brunello^-/-^ tumours (Figure 3B). Additional genes with lesser known or less direct influence on this pathway were also identified in this screen (*GALNT8* (47), *PARVB* (48), *SCN1A* (49), *PPP4R2* (50), *RALGAPB* (51) and *KRT7* (52)), strengthening the co-operative link between SMAD4 deficiency and PI3K/Akt/mTOR pathway deregulation in tumourigenesis.

**Figure 3.**
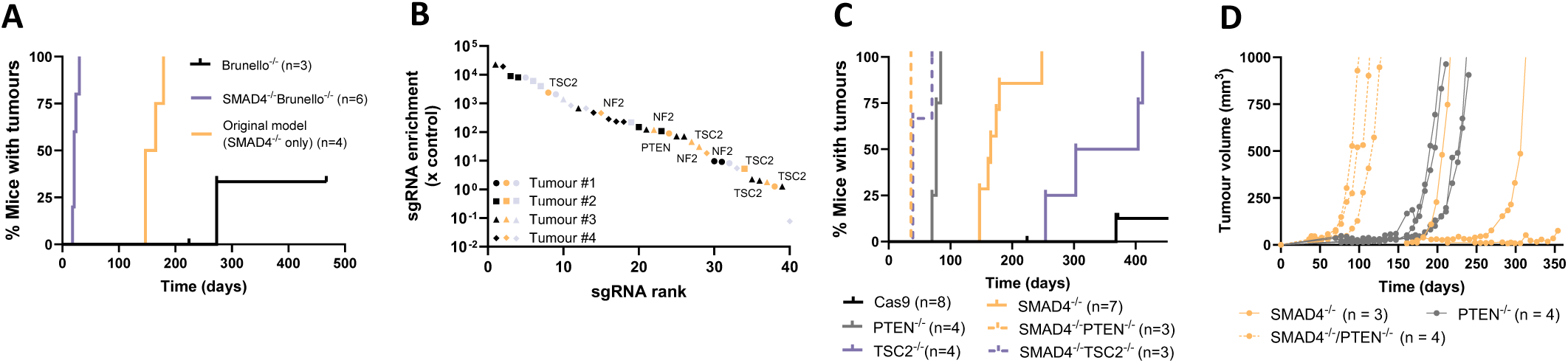
SMAD4 loss and mTOR deregulation co-operate to drive tumourigenesis. (A) Kaplan-Meier plot depicting percentage of mice with tumours over time in CRISPR-Cas9 knockout screen. (B) Scatter plot showing enrichment of individual sgRNAs as identified in tumours from tumourigenesis screen. Orange symbols represent canonical regulators of PI3K/Akt/mTOR signalling and grey symbols represent secondary regulators. (C) Kaplan-Meier plot depicting percentage of mice with tumours over time in single sgRNA validation experiments. (D) Plot showing growth rate of tumours in single sgRNA validation experiments.

To validate these findings, *PTEN*^-/-^ and TSC2^-/-^ populations were generated in wildtype SMAD4 and *SMAD4*^-/-^ cells (Figure 3C, Figure S4A), which were then injected into new mice. Both *PTEN*^-/-^ and *TSC2*^-/-^ promoted rapid tumourigenesis of *SMAD4*^-/-^ cells and were less potently tumourigenic on a wildtype SMAD4 background (Figure 3C). Interestingly, whilst *PTEN*^-/-^ only cells rapidly produced palpable tumours, these neoplasms grew very slowly for about 80 days, remaining relatively dormant until approximately day 150, after which time growth rate drastically increased (Figure 3D). This was in stark contrast to the combined *SMAD4*^-/-^*PTEN*^-/-^ cells, which grew quickly immediately after initiation, supporting the notion of a co-operative effect of loss of these two tumour suppressors. We also validated these findings in two patient-derived organoid (PDO) models of Barrett’s oesophagus. We found that dual *SMAD4/PTEN*^-/-^ PDOs exhibited a dramatically faster growth rate than wildtype PDOs, with development of very large, irregular-shaped organoids (Figure S4B). These data additionally suggest a co-operative effect between loss of SMAD4 and deregulation of the PI3K/Akt/mTOR pathway in promoting tumour formation.

### SMAD4-deficient cells and tumours upregulate mTOR signalling and cap-dependent translation initiation

We next aimed to dissect the mechanism underlying this interplay between SMAD4 and the PI3K/Akt/mTOR signalling pathway (Figure 4A). To understand the status of mTOR signalling in *SMAD4* deficient cells, reverse phase protein array (RPPA) analysis was performed on a panel of mTOR-related proteins in cells with either *SMAD4*^-/-^, *PTEN*^-/-^, or both *SMAD4*^-/-^*PTEN*^-/-^ (Figure 4B). Parental cells displayed generally lower phosphorylation of Akt (S473, T308), mTOR (S2448, S2481), and downstream targets p70 S6 Kinase (p70S6K) and RPS6. *PTEN*^-/-^ cells showed a modest increase in phosphorylation of several mTOR pathway components relative to parental cells, such as p-Akt (S473, T308), p-p70S6K (T389, P421, S424) and p-RPS6 (S235, S236, S240, S244). In contrast, *SMAD4*^-/-^ cells exhibited broadly elevated expression across multiple nodes of the pathway, including MYC, mTOR phospho-sites and p70S6K, suggesting stronger pathway activation. Analyses of TCGA data revealed strong negative correlations between SMAD4 protein level and levels of PI3K/Akt/mTOR family proteins and phospho-proteins in patient OAC samples (Figure 4C), supporting our model. The dual *SMAD4*^-/-^*PTEN*^-/-^ cells showed a mixed pattern, with some targets resembling the high-expression profile of *SMAD4*^-/-^ cells (e.g., RPS6 phosphorylation) and others more closely aligning with the lower expression profile of *PTEN*^-/-^ cells. A similar trend was identified in *TSC2*^-/-^ and *SMAD4*^-/-^*TSC2*^-/-^ cells (Figure S5A). Together, these data point to augmented mTOR pathway activity in *SMAD4*^-/-^ cells, and a combined effect in *SMAD4*^-/-^*PTEN*^-/-^ or *SMAD4*^-/-^*TSC2*^-/-^ dual knockout populations.

**Figure 4.**
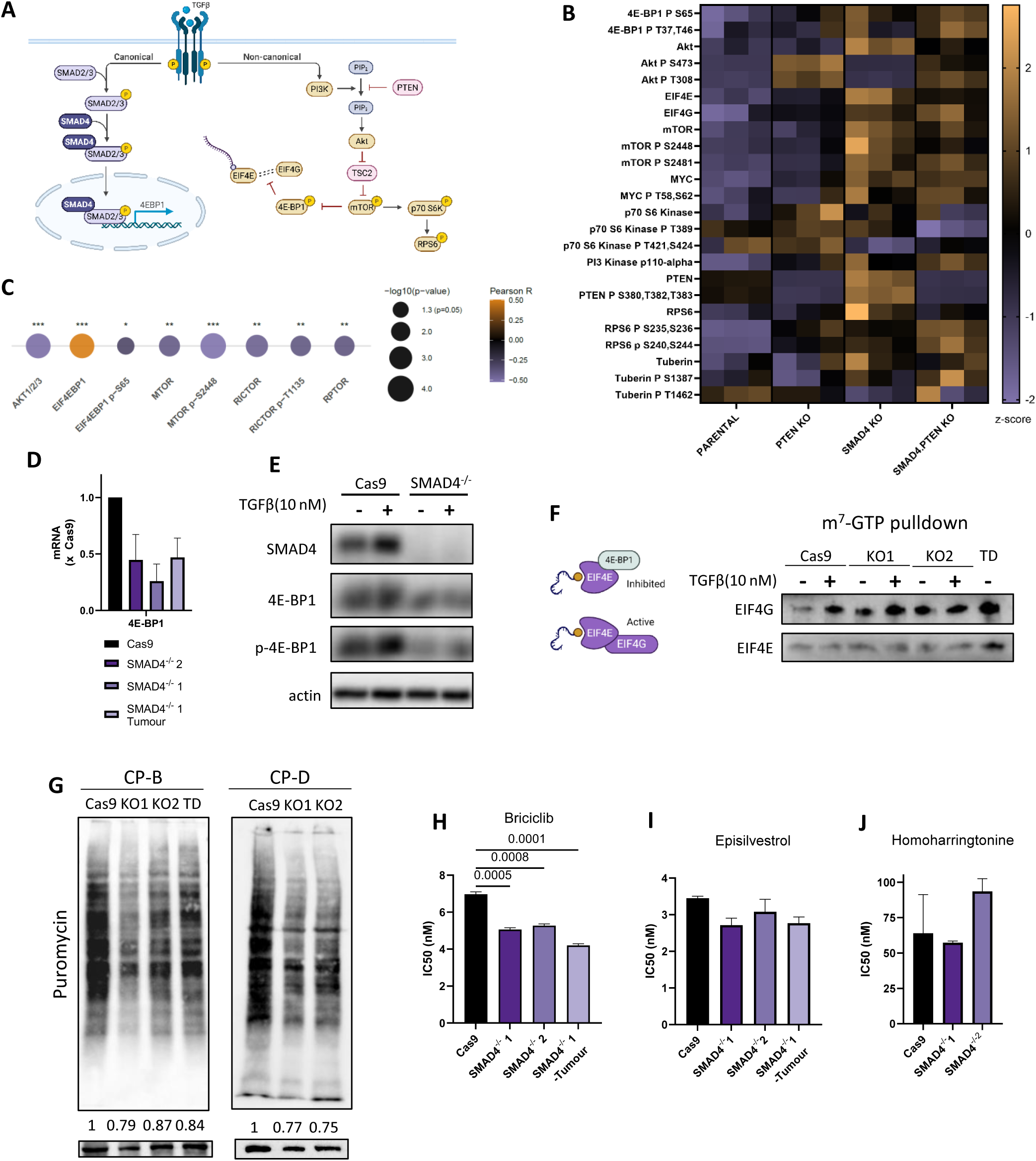
SMAD4-deficient cells and tumours upregulate mTOR signalling and cap-dependent translation initiation. (A) Schematic depicting intersection of TGFβ and PI3K/Akt/mTOR pathways. (B) Heat map depicting protein and phospho-protein expression in CP-B cells as determined by RPPA, depicted as z-scores. (C) Bubble plot showing correlations between SMAD4 protein and PI3K/Akt/mTOR pathway proteins and phospho-proteins in TCGA OAC patient samples. (D) Histogram depicting 4E-BP1 mRNA levels in CP-B cells (n = 3). Error bars represent SEM. (E) Representative immunoblot showing SMAD4, 4E-BP1 and p-4E-BP1 protein levels in CP-B cells in response to 6 h exposure to TGFβ (10 nM) (n = 2). Actin was used as a loading control. (F) (Left) Simplified schematic of eIF4E activated and inhibited states and (right) immunoblot of m7-GTP cap-binding assay showing eIF4G and eIF4E proteins bound to m7-GTP resin (n = 3). KO1/2 = *SMAD4^-/-^* 1/2, TD = *SMAD4^-/-^* tumour-derived. (G) Representative immunoblot showing puromycin incorporation into cell lines in SuNSET assay (n = 2). Histograms depicting IC50 of (H) Briciclib, (I) Episilvestrol and (J) Homoharringtonine in cell lines (n=3).

Of particular note was the impact of the gene perturbations on components of the cap-dependent translation initiation machinery. In the scanning model of translation initiation, eIF4E binds to a m^7^GpppX motif at the 5’ end of mRNA (5’ cap) and associates with eIF4G to form the eIF4F complex (53, 54). The formation of eIF4F is inhibited by 4E-BP1 binding to eIF4E, an association that is inhibited by mTOR mediated phosphorylation of 4E-BP1 (55). We observed increased phosphorylation of 4E-BP1 (S65, T37, T46) in *SMAD4*^-/-^ cells which was amplified further in dual knockout cells (Figure 4B, Figure S5A). Combined with the increased total eIF4E and eIF4G protein observed in these cells, this may demonstrate greater availability of eIF4E to associate with eIF4G to induce cap-dependent translation. Quantitative reverse transcription polymerase chain reaction (qRT-PCR) revealed a trend towards decreased expression of 4E-BP1 mRNA in *SMAD4*^-/-^ cells and tumours derived from *SMAD4*^-/-^ cells (Figure 4D). It was also found that treating parental CP-B cells with TGFβ induced phosphorylation of 4E-BP1 (Figure 4E), in keeping with previous reports of activation of mTOR by a non-canonical TGFβ cascade (56). Moreover, whilst TGFβ appeared to induce phosphorylation of 4E-BP1 in *SMAD4*^-/-^ cells, overall levels of 4E-BP1 were decreased in these cells (Figure 4E). Given that phosphorylated 4E-BP1 is unable to bind to and thereby inhibit eIF4E, these data allude to an increase in available eIF4E in *SMAD4*^-/-^ cells, particularly in the presence of TGFβ.

Thus, a cap-binding pull-down assay was implemented to verify whether eIF4E activity (as well as expression level) was upregulated in *SMAD4*^-/-^ cells. As expected, parental cells showed an increase in eIF4G binding to the m7-GTP cap-bound eIF4E in response to TGFβ (Figure 4F). Furthermore, *SMAD4*^-/-^ cells had higher basal levels of eIF4G binding, which was further increased upon TGFβ stimulation. A SUrface SEnsing of Translation (SUnSET) puromycin incorporation assay (57) confirmed a decrease in global translation rates in both CP-B and CP-D cells without SMAD4, with preferential translation of transcripts encoding only low-to-medium molecular weight proteins (Figure 4G). In line with this interpretation, our multi-omics analyses identified mRNA processing and translation as key clusters of deregulation in SMAD4-deficient cells (Figure 1E). In addition to mitosis-related genes, our genome-wide CRISPR-Cas9 screen identified multiple genes related to translation, including EIF4E, as synthetic lethal in the context of SMAD4 deficiency (Figure S5B). To interrogate this further, we treated cells with a range of translation inhibitors: Briciclib (eIF4E inhibitor, required for cap-dependent translation), Episilvestrol (EIF4A inhibitor, required for global translation) and Homoharringtonine (translation elongation inhibitor). We found increased sensitivity to only Briciclib in *SMAD4*^-/-^ cells (Figure 4H-J, Figure S5C-E), suggesting a specific dependency of SMAD4-deficient cells on cap-dependent translation initiation but not global translation or elongation. These data point to an increase in cap-dependent translation initiation of specific mRNAs in SMAD4-deficient cells, but without a concomitant increase in global translation.

### Translational reprogramming in SMAD4-deficient cells results in post-transcriptional repression of CDK11B

As these data suggested possible reprogramming of mRNA translation in our models, we next investigated the translational profiles of SMAD4-deficient cells via polysome analyses (Figure 5A-B). Here, CP-B parental, *SMAD4*^-/-^ and *SMAD4*^-/-^ tumour-derived cells were lysed and fractionated on a sucrose density gradient to isolate polysome-associated mRNAs. Changes in total mRNA abundance and polysome-bound mRNAs were analysed via RNAseq. This revealed an altered global translational landscape in SMAD4-deficient cells (Figure 5C). SMAD4-deficient cells displayed an increase in free 60S subunits and 80S monosomes, with a concurrent decrease in low-order polysomes (peaks labelled 2-5) and enrichment of high-order polysomes (peaks labelled 7-8). This suggests that, following loss of SMAD4, cells have selectively altered ribosome occupancy on specific subsets of mRNAs, resulting in higher polysome associations on some transcripts at the expense of others. This could represent either increased translation or ribosome stalling on these mRNAs (58). Although the *SMAD4*^-/-^ cell lines displayed enrichment of higher order polysomes, the tumour-derived cell line exhibited a modest reduction in 7-8 ribosome peaks. This pattern is consistent with highly efficient translation (58) and a translatome biased towards shorter, growth-related mRNAs (59). Consistent with this, the *SMAD4*^-/-^ tumour-derived cell line also displayed increased abundance of ribosome and translation-related mRNAs, including EIF4A2, RPL7, RPL8, RPL9, RPL21, RPL23, RPL23A, RPL34, RPL39L, RPS3A, and RPS20 (Figure 5D). Together, these data suggest an association between SMAD4 loss and translational reprogramming with differential ribosome density on a subset of transcripts, and subsequent further changes during transformation to restore efficient translation.

**Figure 5.**
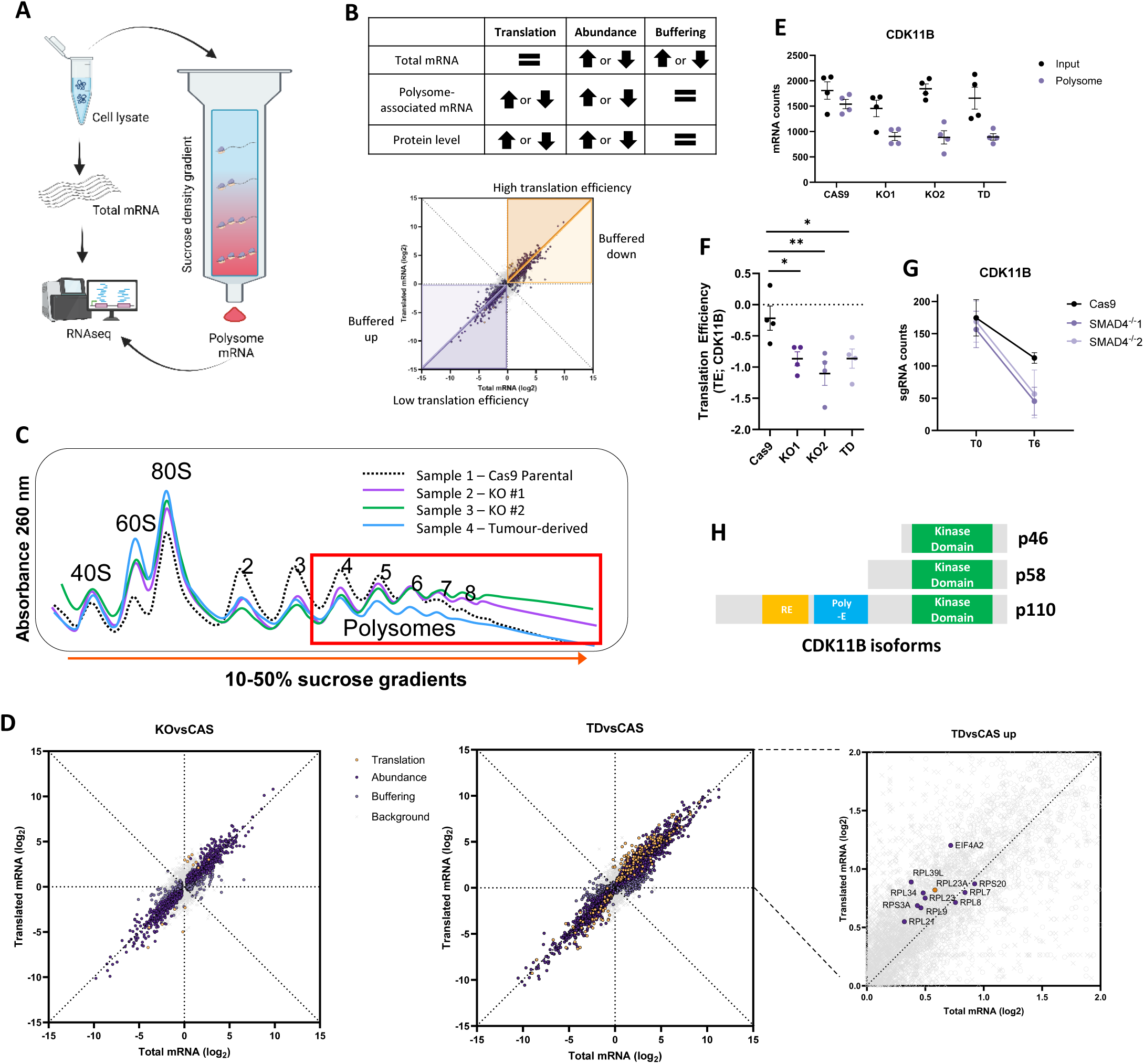
Translational reprogramming in SMAD4-deficient cells results in post-transcriptional repression of CDK11B. Schematics depicting (A) workflow of polysome profiling and (B) interpretation of polysome-sequencing. (C) Polysome traces CP-B cells treated on a 10–40% sucrose gradient (representative of *n*□=□4 biologically independent experiments). (D) Scatterplots showing Log2 fold change (log2FC) total mRNA vs polysome-bound (translated) mRNA in CP-B cells (*n*□=□4), (left) *SMAD4*^-/-^ (KO) versus Cas9 parental (CAS), (centre) *SMAD4*^-/-^ tumour derived (TD) versus Cas9 parental (CAS) and (right) upper right quadrant of TDvsCAS blot showing upregulated mTOR-sensitive mRNA. (E) Plots depicting mRNA counts of whole cell and polysome-bound CDK11B mRNA and (F) translation efficiency of CDK11B in CP-B cells. (G) Raw sgRNA counts for CDK11B in CP-B cells at day 0 (T0) and 6 (T6) in genome-wide CRISPR-Cas9 screen. (H) Schematic illustrating protein isoforms of CDK11B.

Next, the polysome-seq was analysed to detect specific mRNAs that were differentially translated in SMAD4-deficient cells, with a particular focus on mitosis-related genes. We identified that the CDK11B transcript, of which the whole cell mRNA levels remained unchanged across cell lines, was decreased in polysome fractions in *SMAD4*^-/-^ and tumour-derived cells (Figure 5E). Indeed, the translation efficiency quotients for this gene were significantly lower in *SMAD4*^-/-^ and tumour-derived cells (Figure 5F), indicating a reduction in polysome-bound CDK11B mRNA compared to that in the cytoplasmic lysate. Interestingly, our genome-wide CRISPR-Cas9 screen revealed dependency of *SMAD4^-/-^* cells on CDK11B (Figure 5G). The tight translational control of CDK11B has been well described (60), whereby the CDK11B transcript gives rise to multiple protein isoforms (Figure 5H). For most of the cell cycle, CDK11B mRNA produces its full-length 110 kDa protein by way of cap-dependent translation, whilst an additional isoform (p46) is translated from mRNA alternative splicing. However, a third isoform, p58, is translated only during mitosis using an internal start codon (60), and is essential for progression of correct mitosis (61, 62). These data therefore indicate disrupted translational control of CDK11B in SMAD4-deficient cells and tumours.

### Re-introduction of p58 isoform of CDK11B rescues mitosis defects

Given its multiple known roles in maintaining mitotic integrity, we finally aimed to determine whether our observed reduction in CDK11B protein synthesis was influencing the defective mitosis phenotype seen in *SMAD4*^-/-^ cells. To do this, we ectopically expressed cDNA encoding the short, p58 isoform of CDK11B in CP-B cells with or without SMAD4 (Figure 6A) and conducted live-cell imaging to visualise mitoses. Re-expression of CDK11B-p58 significantly reduced the rate of mitotic defects in *SMAD4*^-/-^cells (Figure 6B-C), especially the occurrence of lagging chromosomes (Figure 6D) and failed cytokinesis (Figure 6E). Cells containing the CDK11B-p58 construct also exhibited a shorter time from nuclear envelope breakdown (NEB) to anaphase (Figure 6F), indicating a less error-prone passage through mitosis, but did not alter the percentage of multinuclear cells (Figure 6G). Together, these data suggest that lack of CDK11B-p58 protein in *SMAD4*^-/-^ cells is responsible for the observed mitosis defects.

**Figure 6.**
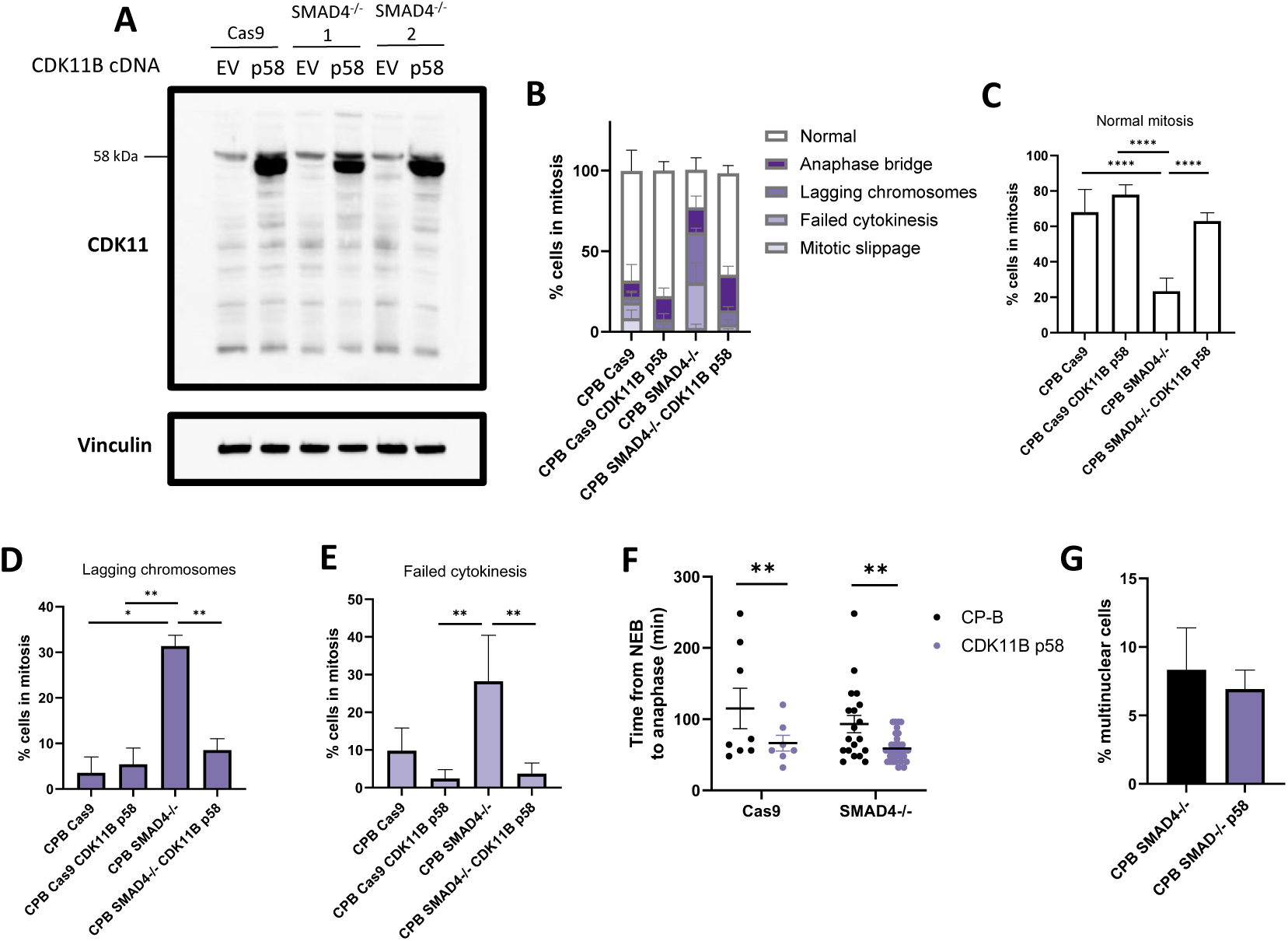
Re-introduction of p58 isoform of CDK11B rescues mitosis defects. (A) Representative immunoblot showing CDK11 protein expression in CP-B cells following ectopic expression of CDK11B-p58 cDNA construct (n = 3). Bar charts showing percentage of (B) mitotic cells with mitosis defects (range 9-38 cells in mitosis visualised per cell line), percentage of cells with (C) normal mitoses, (D) lagging chromosomes, and (E) failed cytokinesis. (F) Plot showing time from nuclear envelope breakdown (NEB) to anaphase in CP-B cells following ectopic expression of CDK11B-p58 cDNA construct. (G) Bar chart showing percentage of multinuclear cells detected via live cell imaging of CP-B cells following ectopic expression of CDK11B-p58 cDNA construct. NB: “CPB Cas9” and “CPB *SMAD4*^-/-^“ data (i.e. untransduced controls) are repeated from Figure 2.

## Discussion

In this study, we delineate a previously undescribed role for SMAD4 in chromosomal instability and tumourigenesis. We provide evidence of SMAD4-associated CIN in multiple patient cohorts, and a deregulated mitosis signature associated with SMAD4 deficiency in our models of p53-mutant pre-cancerous Barrett’s oesophagus. Early Barrett’s metaplasia is marked by low levels of chromosome alterations as compared to dysplastic Barrett’s oesophagus (16), a phenotype attributed to the frequent loss of wildtype p53 function in the latter (28). Whilst OAC has a high mutational burden, it is primarily a cancer of copy number drivers (27, 63–65) and may in fact be characterised by a deregulated mitotic spindle signature (41). The role of p53 in suppressing CIN has been extensively examined, but mutant p53 is frequently detected in pre-malignant Barrett’s tissue long before progression to invasive cancer (66). Therefore, it appears that loss of p53 is unlikely to be the only factor at play in CIN-driven tumourigenesis, and is insufficient to induce oncogenesis on its own in this disease. Strikingly, in our analysis of patient samples, *TP53*-altered tumours exhibit no significant increase in the proportion of the genome altered compared to wild type *TP53* tumours, except in combination with SMAD4 loss (Figure 1A). A similar effect was observed in multiple gastrointestinal cancers (Figure 1A-C), suggesting the role of SMAD4 in maintaining genome stability is not exclusive to OAC. Interestingly, all but one sample with a SMAD4 alteration was shown to also have either a *TP53* alteration or MDM2 amplification (Table S1). We suggest that loss of p53 function is necessary for a concurrent SMAD4 alteration, potentially due to an unstable genome requiring defective error-sensing mechanisms to allow cell survival in this state. A recent study of mitotic spindle dynamics through OAC progression also appears to support this (67). Scott *et al.* demonstrated increasing mitosis defects across a panel of oesophageal cell lines, from low-grade dysplastic Barrett’s oesophagus through to OAC, and determined that these effects are due to loss of wildtype p53 function. However, the authors did not examine the striking negative association between SMAD4 protein level and observed spindle aberrations in these cell lines that may also explain this phenotype, as shown in our data. This also suggests a protective effect of SMAD4 wild type protein against CIN in mutant p53 cells.

Our combined ‘omics approach supported the notion of SMAD4 maintaining chromosomal stability in this context. Specifically, our transcriptomic and proteomic data revealed that loss of SMAD4 resulted in perturbation of critical regulation of the mitotic spindle and chromosome segregation (Figure 1F-G). This expression pattern is similar to that described by Xu *et al.* in their comparative analysis of oesophageal cancer versus normal oesophageal tissues (41). Whilst Xu *et al.* identified defective mitotic spindle regulation as pertinent in oesophageal cancer, here we propose that SMAD4 loss may be one key cause of this deregulation. Furthering this, there is a notable overlap between differentially expressed genes in our SMAD4-deficiency signature and genes known to be involved in mitotic spindle function and CIN (Figure S2G) (39, 40). Despite no documented evidence of such genes as direct SMAD4 transcriptional targets, our finding that expression of these genes positively correlated with SMAD4 protein levels in patient samples (Figure 1I) and cancer cell lines (Figure S2E-F) suggests that this transcriptional control is common in OAC, and potentially other cancer types.

We detected a significant increase in mitotic errors in SMAD4-deficient cells (Figure 2D-E) and propose that accumulation of such errors may result in accrual of tumour-promoting gene alterations. The stochastic nature of these errors may explain the lag time before tumour initiation in *SMAD4*^-/-^Barrett’s cells (18), as not all errors will result in beneficial gene alterations. Supporting this, our genome-wide tumourigenesis screen identified multiple co-operative drivers that hasten transformation of *SMAD4*^-/-^ Barrett’s cells (Figure 3B-C). Many of these drivers converge on the PI3K/Akt/mTOR pathway, suggesting a strong compound effect of mTOR hyperactivation with SMAD4 loss. Additionally, we propose that the pool of co-operative drivers in SMAD4-deficient tumours may be large, where multiple gene alterations may have effects that co-operate with SMAD4 loss to promote tumour growth. This may be why there are very few significantly co-occurring gene alterations in *SMAD4*-altered tumours (27), but instead a large panel of potential co-operators, including PI3K/Akt/mTOR.

We investigated the cross-reactivity between SMAD4 loss and the mTOR pathway, showing that SMAD4-deficient cells upregulate mTOR signalling, and this is increased further when combined with loss of PTEN or TSC2 (Figure 4B, Figure S5A). Known links between SMAD4/TGFβ signalling and mTOR converge on 4E-BP1 (56), pointing to an effect on cap-dependent translation. In line with this, we see an increase in cap-dependent translation in *SMAD4*^-/-^ cells (Figure 4F) and a concomitant reprogramming of global translation (Figure 5C). Our SuNSET assay demonstrates a decrease in puromycin incorporation in SMAD4-deficient cells (Figure 4G), suggesting delayed elongation or premature termination of peptides in these cells. These data likely represent an initiation-elongation imbalance, where slowed elongation is outpaced by enhanced cap-dependent initiation. Our data suggest that post-transformation, SMAD4-deficient tumour cells may alleviate this initiation-induced elongation stress. It is possible that these cells restore translational capacity through accumulation of an additional mTOR-related gene alteration and subsequent upregulation of mTOR-sensitive mRNAs encoding ribosomal proteins and elongation factors (68). This may explain the faster tumour initiation observed in these cells than untransformed, *SMAD4*^-/-^ cells. In the case of untransformed cells, the observed mTOR-driven increase in cap-dependent initiation may influence ribosome density on specific transcripts at the expense of others. We report that translation of CDK11B, a well-described example of translationally controlled protein expression, is altered in *SMAD4*^-/-^ cells (Figure 5E-F). Whilst whole cell mRNA levels remain the same, ribosome-bound CDK11B transcripts are decreased in SMAD4-deficient tumours. The mitotic index of parental versus *SMAD4*^-/-^ cells is similar (Figure S3B), suggesting that a difference in CDK11B protein levels is unlikely due to an imbalance in cell cycle stage. Given our model depicts upregulation of cap-dependent translation initiation in *SMAD4*^-/-^cells, we propose that alternative translation mechanisms are suppressed in these cells resulting in lower expression of CDK11B-p58.

Finally, we show that reintroduction of CDK11B-p58 is sufficient to rescue multiple mitotic defects in *SMAD4*^-/-^ cells, including lagging chromosomes and failed cytokinesis. This is consistent with previous data showing that CDK11-p58 is essential for maintaining cohesion of sister chromatids, as well as recruitment of ESCRT-III complex and abscission during cytokinesis (61, 62). Lagging chromosomes and errors in chromosome segregation can result in formation of micronuclei (69, 70), which are abundant in our *SMAD4*^-/-^ cells. Interestingly, re-expression of CDK11B-p58 did not significantly reduce the percentage of cells with multiple nuclei in our models. A recent study found that TGFBR2-deficient cells became enriched in cell populations with micronuclei (71), suggesting a protective effect of loss of TGFβ signalling in cell survival with micronuclei. We propose that such complex interplay between multiple signalling pathways may result in perpetuation of multinuclear cells despite re-introduction of CDK11B-p58 alleviating the original cause of these abnormalities.

In sum, our data present a model for SMAD4 as a gatekeeper for CIN through regulation of mitosis via translational control. We demonstrate that SMAD4 loss is associated with mitosis defects, CIN, upregulation of PI3K/Akt/mTOR signalling and altered mRNA translation. A product of this translational reprogramming is decreased expression of CDK11B, which is essential for progression of correct mitosis. Importantly, we show that re-introduction of the CDK11B-p58 isoform is sufficient to reduce mitotic errors in SMAD4-deficient cells, plausibly diminishing their tumourigenic potential. However, there remains a delicate balance between tumour-promoting CIN and lethal volatility of the genome, as excessive genome disruption can result in mitotic catastrophe and cell death (3, 72). Further, cells displaying high levels of CIN are acutely sensitive to inhibitors of mitosis and the DNA damage response (DDR) (73, 74). Mounting evidence suggests that tumour progression can be driven by CIN and ploidy abnormalities that promote evolution but may also predispose cells to sensitivity to further mitotic insults. Through our CRISPR-Cas9 dependency screen, we reveal that *SMAD4*^-/-^ cells are indeed more susceptible to loss of other mitosis regulatory genes. Taken together, these data indicate a new role for SMAD4 in maintaining chromosomal stability in pre-cancerous lesions and unveil potential therapeutic avenues for SMAD4-deficient cancers.

## Materials and methods

### Experimental models

#### Cell lines

The human telomerase reverse transcriptase (hTERT) immortalized Barrett’s epithelial cell lines, CP-B (CP-52731 RRID: CVCL_C452) and CP-D (CP-18821, RRID: CVCL_C454), were obtained from Professor Peter Rabinovitch (University of Washington, Seattle, WA) and cultured in MCDB-153 medium supplemented with 400 ng/mL hydrocortisone, 20 ng/mL epidermal growth factor (EGF), 20 mg/mL adenine, 84 mg/mL bovine pituitary extract (BPE), ITS Liquid Media Supplement (100X) for final concentrations of 2.5 mg/mL insulin, 1.4 mg/mL transferrin, 1.3 ng/mL selenium (all from Sigma-Aldrich, Sydney, Australia), 10 nM cholera toxin, 5% v/v fetal bovine serum (FBS), 375 ng/mL fluconazole (Sigma-Aldrich), and 4 mM L-glutamine (GlutaMAX; Thermo Fisher Scientific, Carlsbad, CA), adjusted to pH 7.2. All cells were maintained at 37°C with 5% CO_2_. Authentication of cell lines was performed via short tandem repeat (STR) analysis with the PowerPlex16 genotyping system (Promega, Madison, WI) and cultures were confirmed to be free from mycoplasma by polymerase chain reaction (PCR) (Cerberus Sciences, Scoresby, VIC). Cells were thawed from early passage stocks and passaged no more than 10 times.

#### Mouse models

NOD-SCID IL-2Rγ^KO^ (NSG) mice were obtained from in-house breeding facility. All animal experiments were approved by the Peter MacCallum Cancer Centre (PMCC) Animal Ethics Committee and undertaken in accordance with the National Health and Medical Research Council (NHMRC) Australian Code of Practice for the Care and Use of Animals for Scientific Purposes.

#### Patient-derived Barrett’s oesophagus organoids

Patients were recruited during hospital visits for either routine surveillance endoscopy or oesophagectomy procedure at Peter MacCallum Cancer Centre (PMCC). Written informed consent was obtained from patients and the collection and use of tissue for this study as approved by the Peter MacCallum Cancer Centre Human Research Ethics Committee (Project #HREC/44873/PMCC-2018). Biopsy specimens were collected from regions of Barrett’s metaplasia in the distal oesophagus using 2.8 mm biopsy forceps. Biopsies were transported on ice in organoid wash media, made with base medium Advanced Dulbecco’s Modified Eagle Medium/F12 (AdMEM/F12; Thermo Fisher Scientific, Waltham, MA) supplemented with 1X GlutaMAX (Thermo Fisher Scientific), 10 mM HEPES (Thermo Fisher Scientific), 2.5 μg/mL Amphotericin B and 0.3 mg/mL Imipenem/Cilastatin. Specimens were processed using a modified protocol, combining previously established methods (75, 76) for optimal conditions. Biopsies were minced using a size 23 scalpel blade and digested with Dulbecco’s Modified Eagle Medium (DMEM; Thermo Fisher Scientific) supplemented with 0.3 mg/mL Imipenem/Cilastatin, 1.5 mg/mL Collagenase II (Thermo Fisher Scientific), 800 U hyaluronidase (Sigma-Aldrich, St. Louis, MO) and 10 μM Y-27632 (ROCK inhibitor; Tocris Bioscience, Bristol, UK). Tissues were incubated in a 37°C water bath for 2-3 h, with regular inversion. The digested suspension was triturated passed through a 70 μm filter and centrifuged for 5 min at 350 x *g* at 4°C. Red blood cells were lysed by adding 2 mL ACK solution (150 mM ammonium chloride (NH_4_Cl; Sigma-Aldrich), 10 mM potassium bicarbonate (KHCO_3_; Sigma-Aldrich) and 100 μM EDTA in water, adjusted to pH 7.2-7.4). After 2 min, 13 mL DMEM with 10% FBS was added and the tube centrifuged again for 5 min at 4°C, the supernatant carefully aspirated and the tube kept on ice. The pellet was resuspended in R&D Systems Cultrex Basement Membrane Extract Type II (BME; In Vitro Technologies, Noble Park, VIC), at approximately 100 μL Cultrex BME for every 1 mm of pellet. Cells were then seeded into 15 μL domes onto pre-warmed (37°C) 48-well flat-bottomed plates, with one dome per well. Plates were inverted to promote dispersal of cells throughout the dome while setting, and incubated at 37°C for 30 min before re-inversion and addition of growth medium (detailed below), including 10 μM Y-27632. All cultures were grown in Intesticult™ Organoid Growth Medium (StemCell Technologies, Vancouver, Canada), supplemented with 1 μg/mL gastrin, 100 ng/mL FGF-10 and 0.25 mg/mL imipenim/cilastatin. During derivation and passage, media was supplemented with 10 μM Y-27632.

### Patient data analysis

For fraction of genome alteration analyses, relevant mRNA and protein expression data as well as clinical attributes were downloaded from publicly available databases. OAC patient samples sourced from The Cancer Genome Atlas (TCGA) Firehose Legacy and Memorial Sloan Kettering (MSK) studies using cBioPortal [https://cbioportal.org] (24, 25). These two studies were selected due to their comprehensive profiling of CNA and mutations. Putative CNAs were determined using the GISTIC 2.0 algorithm. Data were last accessed on 9 August, 2025.

UQ OAC samples were collected from patients that gave written informed consent (HREC/2020/QMS/62117, UQ/2020/HE001913). DNA and RNA were extracted using the Qiagen AllPrep DNA/RNA mini kit according to standard protocol (Qiagen, Germany). DNA was used for Illumina SNP arrays and whole-genome or whole-exome sequencing as previously described (150 bp paired-end; Illumina HiSeq (29, 77). RNA sequencing was performed and processed as previously described (30). These data are accessible through the EGA (https://ega-archive.org/studies/EGAS00001002864). The sequence data are generated from patient samples and therefore are available under restricted access. Data access can be granted via the EGA with completion of an institute data transfer agreement, and data will be available for a defined time period once access has been granted. SNP array data was analysed using ASCAT (version 3.2.0), and HRD scores calculated using published formula (32).

### Generation of knockout cell lines

Top and bottom-strand oligonucleotides were designed with the following overhangs to allow for cloning into the pLentiGuide-Puro vector (Addgene #52963): top strand 5’-CACCG and bottom strand 5’-AAACNNNNNNNNNNNNNNNNNNNNC. Oligos were annealed and amplified and cloned into the vector backbone using Esp3I restriction sites as described previously (78). Cloned plasmids were amplified using NucleoSpin Plasmid Miniprep Kit (Macherey-Nagel, Düren, Germany) as per the manufacturer’s specifications.

For production of lentivirus, 5.5 x 10^5^ HEK293T cells/well were seeded in a 6-well plate and left to adhere overnight at 37°C. Third generation lentivirus packaging mix was created by combining 0.5 μg/μL pMDI, 0.25 μg/μL RSV-REV, 0.3 μg/μL VSV and 1 μg/μL plasmid DNA in neat DMEM (Thermo Fisher Scientific). This mix was first vortexed before adding 9.2 μL polyethylenimine (PEI; Thermo Fisher Scientific) at 1 mg/mL, vortexing again and incubating for 15 min at RT. This mix was then added dropwise to HEK293T cells, which were then incubated at 37°C overnight. After 24 h, the media was removed and replaced with fresh media relative to the target cells (i.e. complete MCDB media for CP-B cells). At 48 h post-transfection, the viral supernatant was collected and passed through a 0.45 μm filter and stored at -80°C.

For lentiviral transduction, 5 x 10^5^ cells/well were seeded in a 12-well plate in 400 μL complete media. One third of the harvested viral stock (approx. 600 μL) was added to each well to a total volume of 1 mL/well plus 8 μg/mL sequabrene. Cells were then centrifuged for 30 min at 1200 x *g* at 30°C and incubated at 37°C for 24 h. Cells were then lifted and transferred to flasks supplemented with 2 μg/mL puromycin (selection marker for pLentiGuide-Puro) and incubated at 37°C for three days to select for successfully transduced cells. sgRNA sequences: *SMAD4* (ATAACAGCTATAACTACAAA), *PTEN* (CCAATTCAGGACCCACACGA), *TSC2* (TCCTTGCGATGTACTCGTCG).

### RNA-sequencing

Total cell RNA was isolated from cells using Nucleospin RNA extraction kit (Macherey-Nagel, Germany), in biological triplicate. RNA concentration and purity were measured with a NanoDrop1000 spectrophotometer (Thermo Fisher Scientific). RNA-sequencing library was prepared with QuantSeq Lexogen 3’mRNA-FW on the Illumina NextSeq 500 system using Single End 75 base reads with a read depth of 3-4 million reads per sample. Sequencing reads were aligned using HISAT2 (79), and gene expression was quantified using HTSeq software (80). Normalised expression was measured in counts-per-million (CPM) on a log_2_ scale, with library size adjustment conducted using the Trimmed Mean of M-values method with edgeR (81) R package. Differential expression analysis was performed using the LIMMA-Voom workflow (82). Gene set enrichment analysis was performed using GSEA. MSigDB gene set collections were used as reference gene sets. Dataset can be found in Table S3.

### Quantitative proteomics

CP-B cells (5.0 × 10^6^) were plated in 15 cm plates and allowed to adhere for 24 h. Cells were washed with ice-cold PBS and lysed at 4°C in RIPA buffer (1 mM EDTA, 1% NP-40, 0.5% sodium deoxycholate, 0.1% SDS, 50 mM sodium fluoride, 1 mM sodium pyrophosphate in PBS) supplemented with protease and phosphatase inhibitors (Roche, Basel, Switzerland). Protein purification, tryptic digest and peptide elution were performed and liquid chromatography tandem mass spectrometry (LC-MS/MS) was conducted, all as previously described (83). Data analysis was performed using MaxQuant (84) and Perseus (85) platforms. Dataset can be found in Table S4.

### CRISPR screening

CP-B cells were engineered to express the Cas9 endonuclease by transduction with the FUCas9Cherry vector (Addgene, Watertown, MA; #70182) and subsequent selection for mCherry-positive cells. Cells were then transduced with a genome-wide sgRNA Brunello (76,441 sgRNAs, Addgene; #73178 (43)), at a MOI of 0.3, aiming for 500-fold starting representation of each guide. Puromycin selection (2 μg/mL) was applied for 3 days and surviving cells were passaged every three days in T175 flasks with 8 × 10^6^ cells per flask for a total of 6 days. 3.2 x 10^7^ cells were used for xenograft transplants. Genomic DNA of cells and xenograft tumours was extracted using the DNeasy Blood and Tissue Kit (Qiagen, Hilden, Germany) and sequencing libraries were generated by PCR as previously detailed (86). The libraries were sequenced on a NextSeq 500 (Illumina, San Diego, CA) to generate 75 base pair single-end reads, which were demultiplexed with Bcl2fastq (v2.17.1.14). The sequencing reads were subsequently trimmed with Cutadapt (v1.7) to remove sgRNA vector derived sequences, then sgRNA reads were counted and the distribution analyzed using MAGeCK (v0.5.8) using read depth (--norm-method total) normalization. MAGeCK score was determined by -log_10_(neg *P*-value) + log_10_(pos *P*-value) and datasets were visualized on GraphPad Prism (v9.0). Dataset can be found in Table S5.

### Metascape analysis

Gene lists including top *SMAD4*^-/-^ synthetic lethal hits (MAGeCK score > 3), RNA-sequencing differentially expressed genes (q-value < 0.05) and proteomics differentially abundant proteins (q-value < 0.05) were entered into the Metascape online resource [https://Metascape.org] (34). Gene lists were analyzed for exact gene matches and overlapping enrichment of the following gene ontology sources: KEGG Pathway, GO Biological Processes, Reactome Gene Sets, Canonical Pathways, Cell Type Signatures, CORUM, TRRUST, DisGeNET, PaGenBase, Transcription Factor Targets, WikiPathways, PANTHER Pathway and COVID. Using all genes in the human genome as the background for enrichment, “enrichment factors” were calculated as the ratio of observed counts to the number of counts expected by chance. Clusters were designated as > 3 similar terms with enrichment factors > 1.5 and p-values < 0.01 (calculated using accumulative hypergeometric distribution).

### Competitive growth assay

Cas9-expressing CP-B cells and two *SMAD4*^-/-^ CP-B clonal populations were transduced with sgRNA targeting *ANAPC11*, and treated with 2 µg/mL puromycin for 72 h. The three cell populations were then pooled and re-seeded at 1.0 x 10^5^ cells/well in 6-well plates and T0 confluency determined utilizing an IncuCyte FLR (Essen BioScience, Ann Arbor, MI) imaging phase and green channels. Media was changed after 3 days and an endpoint reading was taken at T6 (i.e. 6 days post-selection). Percentage of *SMAD4*^-/-^ populations was determined as GFP confluency/total cell confluency. Empty vector control wells were set up and confluency quantified to account for differences in transduction efficiency.

### Drug dosing

For dose response assays, serial dilutions of pharmacological compounds (Prexasertib, SelleckChem S7178; Bricilib, SelleckChem S6533; Episilvestrol, MedChemExpress HY-15359; Homoharringtonine, Sigma SML1091) were added to 96-well plates containing 2 x 10^3^ cells/well. After 120 h incubation, cell viability was determined using AlamarBlue reagent (Life Technologies) and fluorescence was read at 550 nm/590 nm on a Cytation 3 Imaging Reader (BioTek).

### DepMap Analysis

Cell line dependency, mRNA expression and proteomics data were obtained from the cancer Dependency Map (DepMap) portal [https://depmap.org], last accessed on 9 August, 2025. The relevant data were extracted from DepMap Public 22Q2 release of the Cancer Cell Line Encyclopaedia (CCLE) project RNA-sequencing and quantitative proteomics (38). The DepMap platform calculates RNA expression values as transcripts per million (TPM) using the RNA-Seq by Expectation-Maximisation (RSEM) tool and transformed using the formula [log_2_(TPM+1)]. Proteomics on the DepMap portal are obtained from quantitative mass spectrometry of 375 cancer cell lines by the Gygi lab (38).

### Live cell imaging

Cells were seeded at 4 x 10^3^ cells/well in black 96-well glass bottom µ-Plates (Ibidi, Fitchburg, WI) in relevant growth medium and incubated at 37°C with 5% CO_2_ for 72 h to allow for adherence and for cells to reach exponential growth rate. One hour prior to recording, media was replaced to contain 0.75X solution SPY650-DNA dye (Spirochrome, Switzerland). Cells were imaged on Olympus FV3000 (Olympus, Japan) in a controlled chamber at 37°C with 5% CO_2_ for 16-18 h using 60x silicone objective (UPLSAPO-S 60x, NA 1.35, Olympus, Japan). Live cell images were captured in z stacks every 8 min. Transmitted channel was acquired and SPY650-DNA dye was excited by 635nm laser. Image processing was performed using Fiji software (87).

### Nucleus size quantification

For immunofluorescence of cell spheroids, cells were grown on low-attachment plates (Corning, Corning, NY) and allowed to form three-dimensional spheroid structures before suspension in agarose and paraffin embedding. Cross sections of the spheroids were stained with DAPI and coverslipped as above. Slides were imaged with fluorescent microscopy, and the size of nuclei analysed and quantified using Fiji software. Nucleus size was determined by converting images to 8-bit greyscale, applying default colour threshold adjustment uniformly to all images, and analysing particles with a lower limit of 200 pixels^2^ to exclude debris and an upper limit of 1100 pixels^2^ to exclude overlapping nuclei.

### Animal experiments

For cell line xenografts, 5 × 10^6^ cells (8 x 10^6^ for the CRISPR screen) suspended in 100 µL of 1:1 PBS and Matrigel (BD Bioscience, Franklin Lakes, NJ) were subcutaneously injected into the right flank of ∼6 week-old female NSG mice. Tumour volume was assessed with caliper measurements every ∼7 days and calculated using the formula (length × width^2^)/2. At experimental endpoint (tumour volume > 1000 mm^3^), mice were culled, tumours and organs were harvested.

### Immunoblotting

As per established protocols (88), cells were lysed at 4°C in RIPA buffer (1 mM EDTA, 1% (v/v) NP-40, 0.5% (w/v) sodium deoxycholate, 0.1% (w/v) SDS, 50 mM sodium fluoride, 1 mM sodium pyrophosphate in PBS) supplemented with cOmplete Mini protease inhibitor and PhosSTOP phosphatase inhibitor cocktails (Roche, Basel, Switzerland). Equal amounts of protein were boiled, resolved by SDS-PAGE and transferred to PVDF membranes. Membranes were incubated in blocking buffer (5% w/v bovine serum albumin (BSA; Bovogen Biologicals, Keilor East, VIC) in Tris-buffered saline with 0.1% v/v Tween-20 (0.1% TBS-T) for 1 h at RT and probed overnight in primary antibody at 4°C. Blots were washed thrice in 0.1% TBS-T, followed by incubation with peroxidase-conjugated secondary antibody (Dako, Denmark) for 1 h at RT. Protein levels were detected using Clarity Western ECL Substrate (BioRad, Hercules, CA) or ECL Plus Western blotting substrate kit (Thermo Fisher Scientific). Antibodies are detailed in Table S6.

### Reverse phase protein array (RPPA)

Cells were seeded at 1 x 10^6^ cells/well in 6-well plates and allowed to adhere overnight before lysis in CeLyA Lysis Buffer (CLB1; Zeptosens, Bayer, Witterswil, Switzerland), and protein quantified using a Pierce™ Coomassie Plus (Bradford) Protein Assay Kit. Using a Sciclone/Caliper ALH3000 liquid handling robot (Perkin Elmer, Waltham, MA), samples were prepared in 4 dilutions (100%, 63%, 40% and 25%) in 10% CLB1:90% CeLyA Spotting Buffer (CSBL1; Zeptosens) and spotted onto ZeptoChips (Zeptosens) in 3-4 technical replicates using a Nano-plotter-NP2.1 non-contact microarray system (GeSiM, Radeburg, Germany). Chips were blocked for 1 h with blocking buffer (BB1; Zeptosens), incubated with pre-validated primary antibodies (1:500, 20 h), and Alexa Fluor® 647 anti-rabbit secondary antibody (1:1000, 4 h) (Thermo Fisher Scientific). Chips were read on a Zeptosens instrument and software version 3.1 used to calculate the Relative Fluorescence Intensity (RFI). All samples were normalised to the background values reported in the secondary antibody-only negative control. Antibodies are detailed in Table S7.

### Reverse transcription quantitative polymerase chain reaction (RT-qPCR)

Cells were seeded 4 x 10^5^ cells/well in 6-well plates and allowed to adhere overnight. Extraction and purification of RNA was conducted using Nucleospin RNA kit (Macherey–Nagel) before reverse transcription to cDNA using Transcriptor First Strand cDNA Synthesis Kit (Roche) as per manufacturer’s instructions, including the optional denaturing step. Quantitative PCR (qPCR) was conducted using LightCycler 480 SYBR-Green qPCR (Roche) as per manufacturer’s protocol, with gene expression normalised within each sample to GAPDH and analysed using the ΔΔCt method. Primer sequences were obtained using the NCBI Primer Blast application and are detailed in Table S8.

### m7-GTP cap-binding assay

Cells were seeded at 1.0 x 10^6^ cells/well in 6-well plates and allowed to adhere overnight. Cells were treated with 10 nM TGFβ or vehicle (citric acid) for 4 h (or cycloheximide for 6 h) before washing with warm PBS. Cells were then lifted with warm 0.05% trypsin-EDTA and centrifuged at 6000 x *g* for 90 s. The trypsin-EDTA was removed and the pellet washed with warm PBS before centrifuging again at 6000 x *g* for 90 s. The supernatant was removed and cells were lysed in non-denaturing lysis buffer (7 mL dH_2_O, 160 μL 5 M NaCl, 160 μL 1 M Tris-HCl pH 7.4, 40 μL 200 mM NaF, 40 μL 1 M MgCl_2_, 40 μL 1 mM sodium orthovanadate, and 40 μL Igepal) with gentle agitation on ice for 2 h. Cells were centrifuged at 12000 x *g* for 15 min at 4°C to remove cellular debris and the supernatant was retained. Five percent of this supernatant was reserved to be used as the whole cell lysate control and stored at -20°C. The remaining cell lysate was added to a microfuge tube containing pre-washed (with TBS) blank agarose control beads (Thermo Fisher Scientific) in a pre-clearing step to remove proteins that interact with the agarose beads, with gentle agitation on ice for 10 min. The blank beads were pelleted by centrifuging at 500 x *g* for 30 s and the lysate transferred to a microfuge tube containing pre-washed (with TBS) γ-aminophenyl-m^7^GTP agarose C10-linked beads (Jena Bioscience, Jena, Germany). Lysis buffer was then added to wash the blank agarose beads, which were pelleted by centrifugation at 500 x *g* for 30 s and this wash process repeated 5 times before resuspension in lysis buffer and stored at -20°C for use as the non-specific agarose bead binding control. The tube containing cell lysate and γ-aminophenyl-m^7^GTP agarose C10-linked beads was incubated with gentle agitation on ice for 1 h to capture cap-binding proteins. The beads were pelleted by centrifuging at 500 x *g* for 30 s and the supernatant removed and stored at -20°C. Lysis buffer was then added to wash the beads, which were pelleted by centrifugation at 500 x *g* for 30 s and this wash process repeated 5 times before resuspension in lysis buffer and GTP added to a final concentration of 1 mM. The beads + 1 mM GTP was incubated on ice with gentle agitation for 1 h to disassociate proteins that non-specifically interact with m^7^GTP. The beads were then pelleted by centrifuging at 500 x *g* for 30 s and the supernatant removed and stored at -20°C for use as the non-specific GTP-binding control. Lysis buffer was added to wash the beads, which were pelleted by centrifugation at 500 x *g* for 30 s and this wash process repeated 5 times before resuspension in a small volume of lysis buffer and storage at -20°C. This final solution contains the m^7^GTP-bound proteins. All samples and controls were boiled to dissolve the agarose beads prior to immunoblotting as mentioned previously, using 4-20% gradient gels (BioRad).

### SuNSET assay

Cells were seeded 4 x 10^5^ cells/well in 6-well plates and allowed to adhere overnight. Cells were then treated with 100 µg/mL cycloheximide for 2 h and/or 1 µM puromycin for 30 min (57) before harvesting and immunoblotting as mentioned previously.

### Polysome profiling

For polysome profiling, cells were treated with 100□µg/mL cycloheximide for 5□min at 37°C, washed with hypotonic wash buffer (5□mM Tris-HCl (pH 7.5), 2.5□mM MgCl_2_, 1.5□mM KCl, 100□µg/mL cycloheximide) and lysed in a hypotonic lysis buffer (5□mM Tris-HCl (pH 7.5), 2.5□mM MgCl_2_, 1.5□mM KCl, 100□µg/mL cycloheximide, 2□mM DTT, 0.5% Triton X-100, 10 U Promega RNAsin and 0.5% sodium deoxycholate). Lysates were centrifuged at 16,000 x *g* for 7 min at 4°C. 10% of supernatant volume was collected to control for input and mixed with TRIzol reagent (Invitrogen). The remaining sample was loaded onto a 10–50% linear sucrose density gradient and centrifuged at 41,000 RPM using a SW41 Ti rotor (Beckman Coulter) for 1.5□h at 4□°C. Gradients were fractionated (16 fractions per sample) and optical density was continuously recorded at 260□nm using a Triax Flow Cell FC1 (BioComp). Fractions were collected and mixed with an equal volume of TRIzol reagent (Invitrogen). Cytoplasmic and polysome-associated RNA were extracted according to the manufacturer’s instructions and purified using the Qiagen RNeasy Mini Kit (Qiagen 74106). For library preparation, rRNA was depleted using NEBNext® Ultra™ II Directional RNA Library Prep Kit. Libraries were sequenced on the Illumina NextSeq2000 platform.

Coding genes were selected using biomaRt. Anota2seq was used for normalization using the “TMM-log2” option, perform QC analyses, residual outlier tests, building the contrast matrix, select significant genes in the total mRNA and translated mRNA fractions, as well as the identification of translation and buffering. To select genes that were significant for translation, a maximum slope of 2 and a minimum slope of -1 was used, and for genes that were significant for buffering we used a minimum slope of -2 and a maximum slope of 1. An adjusted p-value of 0.05 was used as a cutoff.

### Statistical analyses

All statistical analyses were carried out using GraphPad Prism, Version 9.2 (GraphPad Software Inc, San Diego, CA) or Microsoft Excel, Version 16.56, unless otherwise specified. GI50 dose of single agents was determined by fitting the Hill equation. The difference between data groups was analysed using two-tailed Student’s *t*-test or ANOVA with Tukey’s multiple comparisons post hoc test as indicated in the manuscript. Correlations were determined using Pearson’s R. Significant overlap between large gene sets was determined by calculating representation factor [representation factor = # genes in common / (# in group one x # in group 2) / 20471 genes in genome] and using normal approximation of hypergeometric probability <http://nemates.org/MA/progs/representation.stats.html>. Total number of genes in human genome obtained from <Permalink: http://Apr2022.archive.ensembl.org/Homo_sapiens/Info/Annotation>. Data are represented as mean ± standard error of the mean (SEM) unless otherwise specified. *P*-values < 0.05 were considered as statistically significant.

## Acknowledgements

J.V.M. is a Sustaining Innovation Postdoctoral Research Associate and thanks Astex Pharmaceuticals for funding. E.P.K. was supported by a Prostate Cancer Foundation of Australia Future Leaders Fellowship 2024. The authors acknowledge the Centre for Advanced Histology and Microscopy (RRID:SCR_025432), the Molecular Genomics Core (RRID:SCR_025695), the Victorian Centre for Functional Genomics (RRID:SCR_025582) and the Cancer Research Animal Core at Peter MacCallum Cancer Centre for access and support. The Victorian Centre for Functional Genomics (K.J.S: RRID:SCR_025582) is funded by the Australian Cancer Research Foundation (ACRF), Phenomics Australia (https://ror.org/0201hm243), through funding from the Australian Government’s National Collaborative Research Infrastructure Strategy (NCRIS) program, the Peter MacCallum Cancer Centre Foundation and the University of Melbourne Collaborative Research Infrastructure Program. This work was supported by a National Health and Medical Research Council (NHMRC) Ideas grant (#1182525) awarded to N.J.C.

---

**Supplementary Figure S1 SMAD4 loss is associated with chromosomal instability.** Dot plots showing fraction of the genome altered in TCGA and MSK (A) colorectal, (B) gastric or (C) pancreatic cancer patient samples as a function of alteration in *SMAD4* or *TP53* genes. Error bars represent SEM. (D) Dot plots showing fraction of the genome altered in TCGA and MSK OAC patient samples as a function of alteration of TGFBR2 and MYC genes. Error bars represent SEM. (E) Violin plots indicating median and quartiles of trimmed mean of M values (TMM) for *SMAD4* mRNA expression in SMAD4 deficient or intact samples. (F) Violin plots indicating median and quartiles of chromosomal aberrations in UQ OAC patient samples, stratified by SMAD4 status with “strict” criteria. (G) Violin plots indicating median and quartiles of HRD score factors in UQ OAC patient samples. (H) Violin plots depicting chromosomal aberrations in UQ OAC patient samples with locus 18q21.2 removed. (I) Oncoplot showing distribution of genomic alterations in *SMAD4* and *TP53* in UQ OAC patient samples in “strict” criteria. (A-C) Ordinary one-way ANOVA with Tukey’s multiple comparisons test, (D-H) unpaired Student’s t-test.

**Supplementary Figure S2 SMAD4 loss is associated with a deregulated mitosis signature.** (A) (Top) Network of enriched terms illustrated by nodes, colour-coded by dataset. Blue = RNAseq, red = proteomics, green = synthetic lethal CRISPR. Table lists demarked clusters in node network (1–9). (Right) Circos plot depicting overlap between gene lists from RNA-seq (blue arc), proteomics (red arc) and synthetic lethal (green arc) datasets, where the inner circle (orange) represents the total gene set in each dataset, identical genes between datasets are linked by purple curves and genes in shared enriched ontology terms are linked by light blue curves. (B) Top significantly enriched terms in “cell cycle” cluster. (C) Representative phase contrast image showing CP-B cells present at end point of competitive assay, after transduction with sgANAPC11, where Cas9-only only cells are masked in yellow and SMAD4ko (GFP-expressing) are masked in pink, with quantitative histograms for two different sgRNAs (n = 2). (D) Dose response graph depicting cell viability of CP-B cells after 120 h exposure to indicated doses of Prexasertib, normalised to vehicle, and (right) corresponding IC50 values (n = 3). Scatter plots depicting (E) mRNA expression transcripts per million (TPM) plus 1 and (F) relative protein expression BUB1B, CDCA8, SGO1, NDC80 and FBXO8 as a function of SMAD4 protein expression in pan-cancer cell lines (n = 365). (G) Circos plot depicting overlap between gene lists from RNA-seq (this study) and Rines (2008)(40), Xu (2021)(41) and Carter (2006)(39), where the inner circle represents the gene lists, identical genes are linked by purple curves and genes that occur in shared enriched ontology terms are linked by light blue curves. Error bars represent SEM. Correlation determined by simple linear regression and Pearson’s R.

**Supplementary Figure S3 SMAD4 loss is associated with a deregulated mitosis signature.** (A) Schematic depicting model: CP-B/CP-D Cas9 and *SMAD4*^-/-^ 1 & 2 lines were derived from respective parental cell lines. CP-B *SMAD4*^-/-^ tumour derived cells (TD) were derived from tumours that arose from CP-B *SMAD4*^-/-^ 1 cells that had been xenografted into NSG mice. (B) Bar chart depicting mitotic index of CP-B and CP-D cells with or without SMAD4. Violin plots showing (C) time in mitosis and (D) time from nuclear envelope breakdown (NEB) to anaphase onset. Error bars represent SEM. (B) Unpaired Student’s t-test.

**Supplementary Figure S4 SMAD4 loss and mTOR deregulation co-operate to drive tumourigenesis.** (A) Immunoblot of PTEN or TSC2 in CP-B cells transduced with sgPTEN or sgTSC2, respectively. GAPDH and actin were used as loading controls. (B) Representative brightfield microscope images of two Barrett’s oesophagus organoid cultures (Barrett’s ORG1 and 2) nucleofected with sgRNAs targeting *AAVS1* (sgAAVS1), *SMAD4* (sgSMAD4), *PTEN* (sgPTEN) and *SMAD4* and *PTEN* (sgSMAD4+sgPTEN). Scale bars represent 200 µm.

**Supplementary Figure S5 SMAD4-deficient cells and tumours upregulate mTOR signalling and cap-dependent translation initiation.** (A) Heat map depicting protein and phospho-protein expression in CP-B cells as determined by RPPA, depicted as z-scores. (B) Raw sgRNA counts for EIF4E in CP-B cells at day 0 (T0) and 6 (T6) in genome-wide CRISPR-Cas9 screen. Dose response curves for (C) Briciclib, (D) Episilvestrol and (E) Homoharringtonine in CP-B cells (n=3).

## References

1. Hanahan, D., and Weinberg, R. A. (2011) Hallmarks of cancer: the next generation Cell 144, 646–674 10.1016/j.cell.2011.02.013

2. Hosea, R., Hillary, S., Naqvi, S., Wu, S., and Kasim, V. (2024) The two sides of chromosomal instability: drivers and brakes in cancer Signal Transduct Target Ther 9, 75 10.1038/s41392-024-01767-7

3. Sansregret, L., Vanhaesebroeck, B., and Swanton, C. (2018) Determinants and clinical implications of chromosomal instability in cancer Nat Rev Clin Oncol 15, 139–150 10.1038/nrclinonc.2017.198

4. Kapadia, N., and Nurse, P. (2025) Spatiotemporal orchestration of mitosis by cyclin-dependent kinase Nature 643, 1391–1399 10.1038/s41586-025-09172-y

5. Vasudevan, A., Schukken, K. M., Sausville, E. L., Girish, V., Adebambo, O. A., and Sheltzer, J. M. (2021) Aneuploidy as a promoter and suppressor of malignant growth Nat Rev Cancer 21, 89–103 10.1038/s41568-020-00321-1

6. Drews, R. M., Hernando, B., Tarabichi, M., Haase, K., Lesluyes, T., Smith, P. S. et al. (2022) A pan-cancer compendium of chromosomal instability Nature 606, 976–983 10.1038/s41586-022-04789-9

7. Bennett, A., Bechi, B., Tighe, A., Thompson, S., Procter, D. J., and Taylor, S. S. (2015) Cenp-E inhibitor GSK923295: Novel synthetic route and use as a tool to generate aneuploidy Oncotarget 6, 20921–20932 10.18632/oncotarget.4879

8. Trakala, M., Aggarwal, M., Sniffen, C., Zasadil, L., Carroll, A., Ma, D. et al. (2021) Clonal selection of stable aneuploidies in progenitor cells drives high-prevalence tumorigenesis Genes Dev 35, 1079–1092 10.1101/gad.348341.121

9. Bach, D. H., Zhang, W., and Sood, A. K. (2019) Chromosomal Instability in Tumor Initiation and Development Cancer Res 79, 3995–4002 10.1158/0008-5472.CAN-18-3235

10. Lukow, D. A., Sausville, E. L., Suri, P., Chunduri, N. K., Wieland, A., Leu, J. et al. (2021) Chromosomal instability accelerates the evolution of resistance to anti-cancer therapies Dev Cell 56, 2427–2439 e2424 10.1016/j.devcel.2021.07.009

11. Salgueiro, L., Buccitelli, C., Rowald, K., Somogyi, K., Kandala, S., Korbel, J. O., et al. (2020) Acquisition of chromosome instability is a mechanism to evade oncogene addiction EMBO Mol Med 12, e10941 10.15252/emmm.201910941

12. Bass, A. J., Thorsson, V., Shmulevich, I., Reynolds, S. M., Miller, M., Bernard, B. et al. (2014) Comprehensive molecular characterization of gastric adenocarcinoma Nature 513, 202–209 10.1038/nature13480

13. Kim, J., Bowlby, R., Mungall, A. J., Robertson, A. G., Odze, R. D., Cherniack, A. D. et al. (2017) Integrated genomic characterization of oesophageal carcinoma Nature 541, 169–175 10.1038/nature20805

14. Muzny, D. M., Bainbridge, M. N., Chang, K., Dinh, H. H., Drummond, J. A., Fowler, G. et al. (2012) Comprehensive molecular characterization of human colon and rectal cancer Nature 487, 330–337 10.1038/nature11252

15. Nowicki-Osuch, K., Zhuang, L., Jammula, S., Bleaney, C. W., Mahbubani, K. T., Devonshire, G. et al. (2021) Molecular phenotyping reveals the identity of Barrett’s esophagus and its malignant transition Science 373, 760–767 10.1126/science.abd1449

16. Stachler, M. D., Taylor-Weiner, A., Peng, S., McKenna, A., Agoston, A. T., Odze, R. D. et al. (2015) Paired exome analysis of Barrett’s esophagus and adenocarcinoma Nat Genet 47, 1047–1055 10.1038/ng.3343

17. Killcoyne, S., and Fitzgerald, R. C. (2021) Evolution and progression of Barrett’s oesophagus to oesophageal cancer Nat Rev Cancer 21, 731–741 10.1038/s41568-021-00400-x

18. Gotovac, J. R., Kader, T., Milne, J. V., Fujihara, K. M., Lara-Gonzalez, L. E., Gorringe, K. L. et al. (2021) Loss of SMAD4 Is Sufficient to Promote Tumorigenesis in a Model of Dysplastic Barrett’s Esophagus Cell Mol Gastroenterol Hepatol 12, 689–713 10.1016/j.jcmgh.2021.03.008

19. Wang, J. D., Jin, K., Chen, X. Y., Lv, J. Q., and Ji, K. W. (2017) Clinicopathological significance of SMAD4 loss in pancreatic ductal adenocarcinomas: a systematic review and meta-analysis Oncotarget 8, 16704–16711 10.18632/oncotarget.14335

20. Zhao, M., Mishra, L., and Deng, C. X. (2018) The role of TGF-beta/SMAD4 signaling in cancer Int J Biol Sci 14, 111–123 10.7150/ijbs.23230

21. Massague, J. (2008) TGFbeta in Cancer Cell 134, 215–230 10.1016/j.cell.2008.07.001

22. Massague, J. (2012) TGFbeta signalling in context Nat Rev Mol Cell Biol 13, 616–630 10.1038/nrm3434

23. Shi, C., Tao, S., Ren, G., Yang, E. J., Shu, X., Mou, P. K. et al. (2022) Aurora kinase A inhibition induces synthetic lethality in SMAD4-deficient colorectal cancer cells via spindle assembly checkpoint activation Oncogene 10.1038/s41388-022-02293-y

24. Cerami, E., Gao, J., Dogrusoz, U., Gross, B. E., Sumer, S. O., Aksoy, B. A. et al. (2012) The cBio Cancer Genomics Portal: An Open Platform for Exploring Multidimensional Cancer Genomics Data Cancer Discovery 2, 401–404 10.1158/2159-8290.Cd-12-0095

25. Gao, J., Aksoy, B. A., Dogrusoz, U., Dresdner, G., Gross, B., Sumer, S. O., et al. (2013) Integrative analysis of complex cancer genomics and clinical profiles using the cBioPortal Sci Signal 6, pl1 10.1126/scisignal.2004088

26. Sihag, S., Nussenzweig, S. C., Walch, H. S., Hsu, M., Tan, K. S., Sanchez-Vega, F. et al. (2021) Next-Generation Sequencing of 487 Esophageal Adenocarcinomas Reveals Independently Prognostic Genomic Driver Alterations and Pathways Clin Cancer Res 27, 3491–3498 10.1158/1078-0432.Ccr-20-4707

27. Frankell, A. M., Jammula, S., Li, X., Contino, G., Killcoyne, S., Abbas, S. et al. (2019) The landscape of selection in 551 esophageal adenocarcinomas defines genomic biomarkers for the clinic Nat Genet 51, 506–516 10.1038/s41588-018-0331-5

28. Soto, M., Raaijmakers, J. A., Bakker, B., Spierings, D. C. J., Lansdorp, P. M., Foijer, F. et al. (2017) p53 Prohibits Propagation of Chromosome Segregation Errors that Produce Structural Aneuploidies Cell Rep 19, 2423–2431 10.1016/j.celrep.2017.05.055

29. Brosda, S., Aoude, L. G., Bonazzi, V. F., Patel, K., Lonie, J. M., Belle, C. J. et al. (2024) Spatial intra-tumour heterogeneity and treatment-induced genomic evolution in oesophageal adenocarcinoma: implications for prognosis and therapy Genome Med 16, 90 10.1186/s13073-024-01362-z

30. M, M. N., Newell, F., Aoude, L. G., Bonazzi, V. F., Patel, K., Lampe, G., et al. (2023) Multi-omic features of oesophageal adenocarcinoma in patients treated with preoperative neoadjuvant therapy Nat Commun 14, 3155 10.1038/s41467-023-38891-x

31. Van Loo, P., Nordgard, S. H., Lingjærde, O. C., Russnes, H. G., Rye, I. H., Sun, W. et al. (2010) Allele-specific copy number analysis of tumors Proc Natl Acad Sci U S A 107, 16910–16915 10.1073/pnas.1009843107

32. Telli, M. L., Timms, K. M., Reid, J., Hennessy, B., Mills, G. B., Jensen, K. C. et al. (2016) Homologous Recombination Deficiency (HRD) Score Predicts Response to Platinum-Containing Neoadjuvant Chemotherapy in Patients with Triple-Negative Breast Cancer Clin Cancer Res 22, 3764–3773 10.1158/1078-0432.Ccr-15-2477

33. Palanca-Wessels, M. C., Barrett, M. T., Galipeau, P. C., Rohrer, K. L., Reid, B. J., and Rabinovitch, P. S. (1998) Genetic analysis of long-term Barrett’s esophagus epithelial cultures exhibiting cytogenetic and ploidy abnormalities Gastroenterology 114, 295–304 10.1016/s0016-5085(98)70480-9

34. Zhou, Y., Zhou, B., Pache, L., Chang, M., Khodabakhshi, A. H., Tanaseichuk, O. et al. (2019) Metascape provides a biologist-oriented resource for the analysis of systems-level datasets Nat Commun 10, 1523 10.1038/s41467-019-09234-6

35. Peters, J.-M. (2006) The anaphase promoting complex/cyclosome: a machine designed to destroy Nature Reviews Molecular Cell Biology 7, 644–656 10.1038/nrm1988

36. Castro, A., Bernis, C., Vigneron, S., Labbé, J.-C., and Lorca, T. (2005) The anaphase-promoting complex: a key factor in the regulation of cell cycle Oncogene 24, 314–325 10.1038/sj.onc.1207973

37. Cuinat, S., Bézieau, S., Deb, W., Mercier, S., Vignard, V., Isidor, B. et al. (2024) Understanding neurodevelopmental proteasomopathies as new rare disease entities: A review of current concepts, molecular biomarkers, and perspectives Genes Dis 11, 101130 10.1016/j.gendis.2023.101130

38. Nusinow, D. P., Szpyt, J., Ghandi, M., Rose, C. M., McDonald, E. R., III, Kalocsay, M., et al. (2020) Quantitative Proteomics of the Cancer Cell Line Encyclopedia Cell 180, 387–402.e316 10.1016/j.cell.2019.12.023

39. Carter, S. L., Eklund, A. C., Kohane, I. S., Harris, L. N., and Szallasi, Z. (2006) A signature of chromosomal instability inferred from gene expression profiles predicts clinical outcome in multiple human cancers Nat Genet 38, 1043–1048 10.1038/ng1861

40. Rines, D. R., Gomez-Ferreria, M. A., Zhou, Y., DeJesus, P., Grob, S., Batalov, S., et al. (2008) Whole genome functional analysis identifies novel components required for mitotic spindle integrity in human cells Genome Biol 9, R44 10.1186/gb-2008-9-2-r44

41. Xu, W., Xu, J., Wang, Z., and Jiang, Y. (2021) Weighted Gene Correlation Network Analysis Identifies Specific Functional Modules and Genes in Esophageal Cancer J Oncol 2021, 8223263 10.1155/2021/8223263

42. Bloomfield, M., Huth, S. M., McCausland, D. S., Saad, R., Bano, N., Chau, T. N. et al. (2026) Cell and Nuclear Size are Associated with Chromosomal Instability and Tumorigenicity in Cancer Cells that Undergo Whole Genome Doubling Cancer Research 10.1158/0008-5472.Can-24-3718

43. Doench, J. G., Fusi, N., Sullender, M., Hegde, M., Vaimberg, E. W., Donovan, K. F. et al. (2016) Optimized sgRNA design to maximize activity and minimize off-target effects of CRISPR-Cas9 Nat Biotechnol 34, 184–191 10.1038/nbt.3437

44. Lee, Y.-R., Chen, M., and Pandolfi, P. P. (2018) The functions and regulation of the PTEN tumour suppressor: new modes and prospects Nature Reviews Molecular Cell Biology 19, 547–562 10.1038/s41580-018-0015-0

45. Inoki, K., Li, Y., Zhu, T., Wu, J., and Guan, K.-L. (2002) TSC2 is phosphorylated and inhibited by Akt and suppresses mTOR signalling Nature Cell Biology 4, 648–657 10.1038/ncb839

46. James, M. F., Han, S., Polizzano, C., Plotkin, S. R., Manning, B. D., Stemmer-Rachamimov, A. O. et al. (2009) NF2/merlin is a novel negative regulator of mTOR complex 1, and activation of mTORC1 is associated with meningioma and schwannoma growth Mol Cell Biol 29, 4250–4261 10.1128/mcb.01581-08

47. Huang, T., Meng, F., Huang, H., Wang, L., Wang, L., Liu, Y. et al. (2022) GALNT8 suppresses breast cancer cell metastasis potential by regulating EGFR O-GalNAcylation Biochemical and Biophysical Research Communications 601, 16–23 10.1016/j.bbrc.2022.02.072

48. Liu, R., Shang, W., Liu, Y., Xie, Y., Luan, J., Zhang, T. et al. (2024) Inhibition of the ILK-AKT pathway by upregulation of PARVB contributes to the cochlear cell death in Fascin2 gene knockout mice Cell Death Discovery 10, 89 10.1038/s41420-024-01851-5

49. Tsai, C. W., Ho, S. Y., Chen, I. C., Chang, K. C., Chen, H. J., Tsai, F. C. et al. (2025) Abnormal increased mTOR signaling regulates seizure threshold in Dravet syndrome Neuropharmacology 262, 110166 10.1016/j.neuropharm.2024.110166

50. Dong, M. Z., Ouyang, Y. C., Gao, S. C., Gu, L. J., Guo, J. N., Sun, S. M. et al. (2024) Protein phosphatase 4 maintains the survival of primordial follicles by regulating autophagy in oocytes Cell Death Dis 15, 658 10.1038/s41419-024-07051-4

51. Martin, T. D., Chen, X. W., Kaplan, R. E., Saltiel, A. R., Walker, C. L., Reiner, D. J. et al. (2014) Ral and Rheb GTPase activating proteins integrate mTOR and GTPase signaling in aging, autophagy, and tumor cell invasion Mol Cell 53, 209–220 10.1016/j.molcel.2013.12.004

52. Zhao, Z., Meng, M., Yao, J., Zhou, H., Chen, Y., Liu, J. et al. (2023) The long non-coding RNA keratin-7 antisense acts as a new tumor suppressor to inhibit tumorigenesis and enhance apoptosis in lung and breast cancers Cell Death Dis 14, 293 10.1038/s41419-023-05802-3

53. Gross, J. D., Moerke, N. J., von der Haar, T., Lugovskoy, A. A., Sachs, A. B., McCarthy, J. E. G., et al. (2003) Ribosome Loading onto the mRNA Cap Is Driven by Conformational Coupling between eIF4G and eIF4E Cell 115, 739–750 10.1016/S0092-8674(03)00975-9

54. Grifo, J. A., Tahara, S. M., Morgan, M. A., Shatkin, A. J., and Merrick, W. C. (1983) New initiation factor activity required for globin mRNA translation J Biol Chem 258, 5804–5810,

55. Pause, A., Belsham, G. J., Gingras, A. C., Donzé, O., Lin, T. A., Lawrence, J. C., Jr. et al. (1994) Insulin-dependent stimulation of protein synthesis by phosphorylation of a regulator of 5’-cap function Nature 371, 762–767 10.1038/371762a0

56. Azar, R., Alard, A., Susini, C., Bousquet, C., and Pyronnet, S. (2009) 4E-BP1 is a target of Smad4 essential for TGFbeta-mediated inhibition of cell proliferation EMBO J 28, 3514–3522 10.1038/emboj.2009.291

57. Schmidt, E. K., Clavarino, G., Ceppi, M., and Pierre, P. (2009) SUnSET, a nonradioactive method to monitor protein synthesis Nature Methods 6, 275–277 10.1038/nmeth.1314

58. Rodriguez-Martinez, A., and Young-Baird, S. K. (2025) Polysome profiling is an extensible tool for the analysis of bulk protein synthesis, ribosome biogenesis, and the specific steps in translation Mol Biol Cell 36, mr2 10.1091/mbc.E24-08-0341

59. Thoreen, C. C., Chantranupong, L., Keys, H. R., Wang, T., Gray, N. S., and Sabatini, D. M. (2012) A unifying model for mTORC1-mediated regulation of mRNA translation Nature 485, 109–113 10.1038/nature11083

60. Cornelis, S., Bruynooghe, Y., Denecker, G., Van Huffel, S., Tinton, S., and Beyaert, R. (2000) Identification and characterization of a novel cell cycle-regulated internal ribosome entry site Mol Cell 5, 597–605 10.1016/s1097-2765(00)80239-7

61. Hu, D., Valentine, M., Kidd, V. J., and Lahti, J. M. (2007) CDK11p58 is required for the maintenance of sister chromatid cohesion Journal of Cell Science 120, 2424–2434 10.1242/jcs.007963

62. Renshaw, M. J., Panagiotou, T. C., Lavoie, B. D., and Wilde, A. (2019) CDK11(p58)-cyclin L1β regulates abscission site assembly J Biol Chem 294, 18639–18649 10.1074/jbc.RA119.009107

63. Killcoyne, S., Gregson, E., Wedge, D. C., Woodcock, D. J., Eldridge, M. D., de la Rue, R. et al. (2020) Genomic copy number predicts esophageal cancer years before transformation Nat Med 26, 1726–1732 10.1038/s41591-020-1033-y

64. Li, X., Paulson, T. G., Galipeau, P. C., Sanchez, C. A., Liu, K., Kuhner, M. K. et al. (2015) Assessment of Esophageal Adenocarcinoma Risk Using Somatic Chromosome Alterations in Longitudinal Samples in Barrett’s Esophagus Cancer Prev Res (Phila) 8, 845–856 10.1158/1940-6207.CAPR-15-0130

65. Hoppe, S., Jonas, C., Wenzel, M. C., Velazquez Camacho, O., Arolt, C., Zhao, Y., et al. (2021) Genomic and Transcriptomic Characteristics of Esophageal Adenocarcinoma Cancers (Basel) 13, 10.3390/cancers13174300

66. Bansal, A., and Fitzgerald, R. C. (2015) Biomarkers in Barrett’s Esophagus: Role in Diagnosis, Risk Stratification, and Prediction of Response to Therapy Gastroenterology Clinics 44, 373–390 10.1016/j.gtc.2015.02.008

67. Scott, S. J., Li, X., Jammula, S., Devonshire, G., Lindon, C., Fitzgerald, R. C. et al. (2021) Evidence that polyploidy in esophageal adenocarcinoma originates from mitotic slippage caused by defective chromosome attachments Cell Death Differ 28, 2179–2193 10.1038/s41418-021-00745-8

68. Gandin, V., Masvidal, L., Hulea, L., Gravel, S. P., Cargnello, M., McLaughlan, S. et al. (2016) nanoCAGE reveals 5’ UTR features that define specific modes of translation of functionally related MTOR-sensitive mRNAs Genome Res 26, 636–648 10.1101/gr.197566.115

69. Mazzagatti, A., Engel, J. L., and Ly, P. (2024) Boveri and beyond: Chromothripsis and genomic instability from mitotic errors Mol Cell 84, 55–69 10.1016/j.molcel.2023.11.002

70. Hoffelder, D. R., Luo, L., Burke, N. A., Watkins, S. C., Gollin, S. M., and Saunders, W. S. (2004) Resolution of anaphase bridges in cancer cells Chromosoma 112, 389–397 10.1007/s00412-004-0284-6

71. Adams, D. J., Barlas, B., McIntyre, R. E., Salguero, I., van der Weyden, L., Barros, A. et al. (2024) Genetic determinants of micronucleus formation in vivo Nature 627, 130–136 10.1038/s41586-023-07009-0

72. Gronroos, E., and Lopez-Garcia, C. (2018) Tolerance of Chromosomal Instability in Cancer: Mechanisms and Therapeutic Opportunities Cancer Res 78, 6529–6535 10.1158/0008-5472.CAN-18-1958

73. Beksac, M., Balli, S., and Akcora Yildiz, D. (2020) Drug Targeting of Genomic Instability in Multiple Myeloma Front Genet 11, 228 10.3389/fgene.2020.00228

74. Southgate, H. E. D., Chen, L., Curtin, N. J., and Tweddle, D. A. (2020) Targeting the DNA Damage Response for the Treatment of High Risk Neuroblastoma Front Oncol 10, 371 10.3389/fonc.2020.00371

75. Sato, T., Stange, D. E., Ferrante, M., Vries, R. G., Van Es, J. H., Van den Brink, S., et al. (2011) Long-term expansion of epithelial organoids from human colon, adenoma, adenocarcinoma, and Barrett’s epithelium Gastroenterology 141, 1762–1772 10.1053/j.gastro.2011.07.050

76. Li, X., Francies, H. E., Secrier, M., Perner, J., Miremadi, A., Galeano-Dalmau, N. et al. (2018) Organoid cultures recapitulate esophageal adenocarcinoma heterogeneity providing a model for clonality studies and precision therapeutics Nature Communications 9, 2983 10.1038/s41467-018-05190-9

77. Bonazzi, V. F., Aoude, L. G., Brosda, S., Lonie, J. M., Patel, K., Bradford, J. J. et al. (2022) ctDNA as a biomarker of progression in oesophageal adenocarcinoma ESMO Open 7, 100452 10.1016/j.esmoop.2022.100452

78. Sanjana, N. E., Shalem, O., and Zhang, F. (2014) Improved vectors and genome-wide libraries for CRISPR screening Nat Methods 11, 783–784 10.1038/nmeth.3047

79. Kim, D., Langmead, B., and Salzberg, S. L. (2015) HISAT: a fast spliced aligner with low memory requirements Nat Methods 12, 357–360 10.1038/nmeth.3317

80. Anders, S., Pyl, P. T., and Huber, W. (2015) HTSeq--a Python framework to work with high-throughput sequencing data Bioinformatics 31, 166–169 10.1093/bioinformatics/btu638

81. Robinson, M. D., and Oshlack, A. (2010) A scaling normalization method for differential expression analysis of RNA-seq data Genome Biol 11, R25 10.1186/gb-2010-11-3-r25

82. Law, C. W., Chen, Y., Shi, W., and Smyth, G. K. (2014) voom: Precision weights unlock linear model analysis tools for RNA-seq read counts Genome Biol 15, R29 10.1186/gb-2014-15-2-r29

83. Fujihara, K. M., Zhang, B. Z., Jackson, T. D., Ogunkola, M. O., Nijagal, B., Milne, J. V., et al. (2022) Eprenetapopt triggers ferroptosis, inhibits NFS1 cysteine desulfurase, and synergizes with serine and glycine dietary restriction Sci Adv 8, eabm9427 10.1126/sciadv.abm9427

84. Tyanova, S., Temu, T., and Cox, J. (2016) The MaxQuant computational platform for mass spectrometry-based shotgun proteomics Nature Protocols 11, 2301–2319 10.1038/nprot.2016.136

85. Tyanova, S., Temu, T., Sinitcyn, P., Carlson, A., Hein, M. Y., Geiger, T. et al. (2016) The Perseus computational platform for comprehensive analysis of (prote)omics data Nature Methods 13, 731–740 10.1038/nmeth.3901

86. Joung, J., Konermann, S., Gootenberg, J. S., Abudayyeh, O. O., Platt, R. J., Brigham, M. D. et al. (2017) Genome-scale CRISPR-Cas9 knockout and transcriptional activation screening Nat Protoc 12, 828–863 10.1038/nprot.2017.016

87. Schindelin, J., Arganda-Carreras, I., Frise, E., Kaynig, V., Longair, M., Pietzsch, T. et al. (2012) Fiji: an open-source platform for biological-image analysis Nature Methods 9, 676–682 10.1038/nmeth.2019

88. Liu, D. S., Duong, C. P., Haupt, S., Montgomery, K. G., House, C. M., Azar, W. J., et al. (2017) Inhibiting the system xC−/glutathione axis selectively targets cancers with mutant-p53 accumulation Nature Communications 8, 14844 10.1038/ncomms14844

