## Supplementary Figures for "SMAD4 loss drives chromosomal instability during tumourigenesis via translational reprogramming"

**Figure S1**

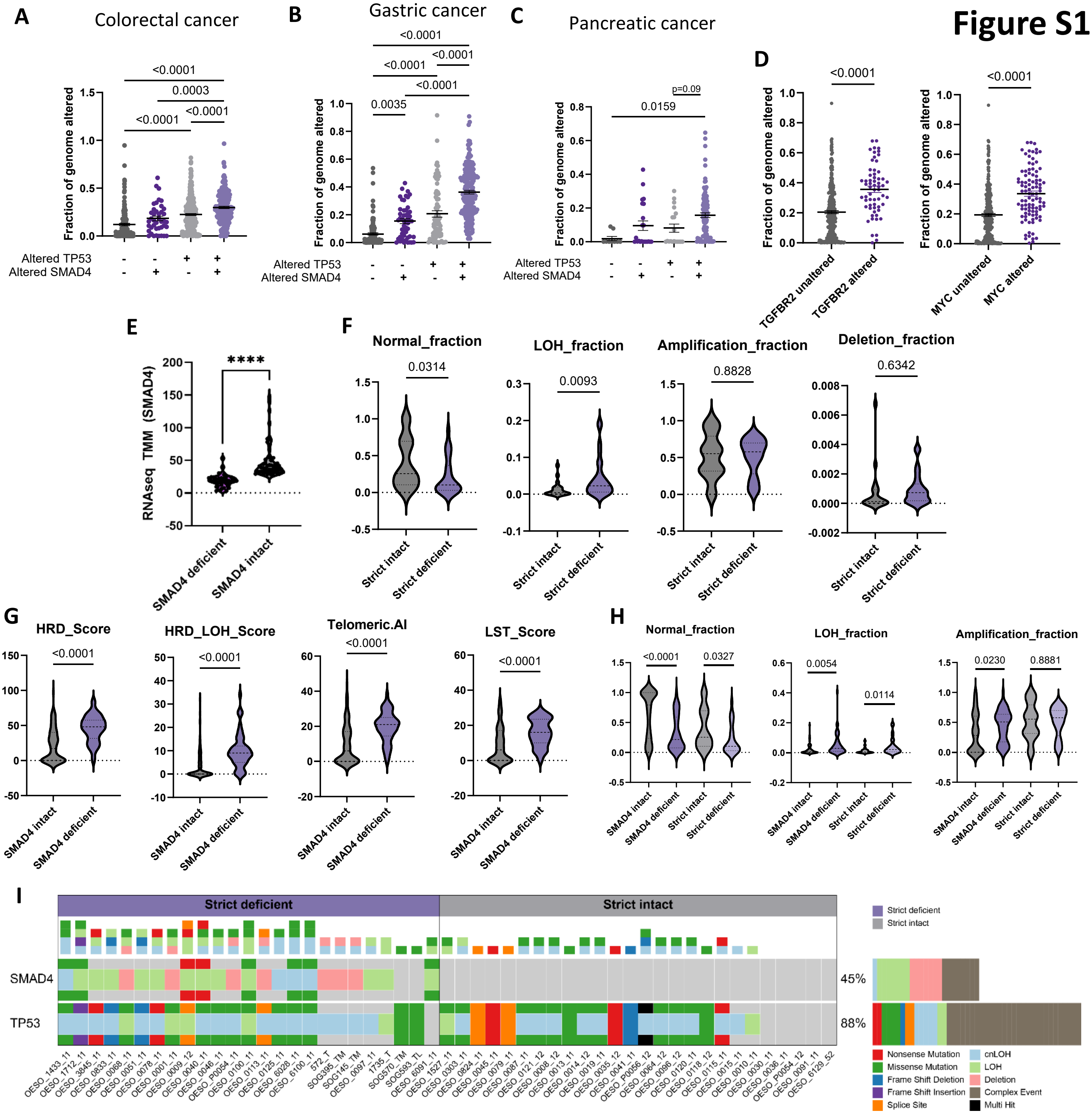

A
Colorectal cancer
Fraction of genome altered
Altered TP53
Altered SMAD4
<0.0001
0.0003
<0.0001
<0.0001
B
Gastric cancer
<0.0001
<0.0001
<0.0001
<0.0001
0.0035
C
Pancreatic cancer
p=0.09
0.0159
D
TGFBR2 unaltered
TGFBR2 altered
MYC unaltered
MYC altered
E
RNAseq TMM (SMAD4)
****
SMAD4 deficient
SMAD4 intact
F
Normal_fraction
0.0314
LOH_fraction
0.0093
Amplification_fraction
0.8828
Deletion_fraction
0.6342
Strict intact
Strict deficient
G
HRD_Score
HRD_LOH_Score
Telomeric.AI
LST_Score
<0.0001
SMAD4 intact
SMAD4 deficient
H
Normal_fraction
<0.0001
0.0327
LOH_fraction
0.0054
0.0114
Amplification_fraction
0.0230
0.8881
I
Strict deficient
Strict intact
SMAD4
45%
TP53
88%
Nonsense Mutation
Missense Mutation
Frame Shift Deletion
Frame Shift Insertion
Splice Site
cnLOH
LOH
Deletion
Complex Event
Multi Hit

### Figure S2

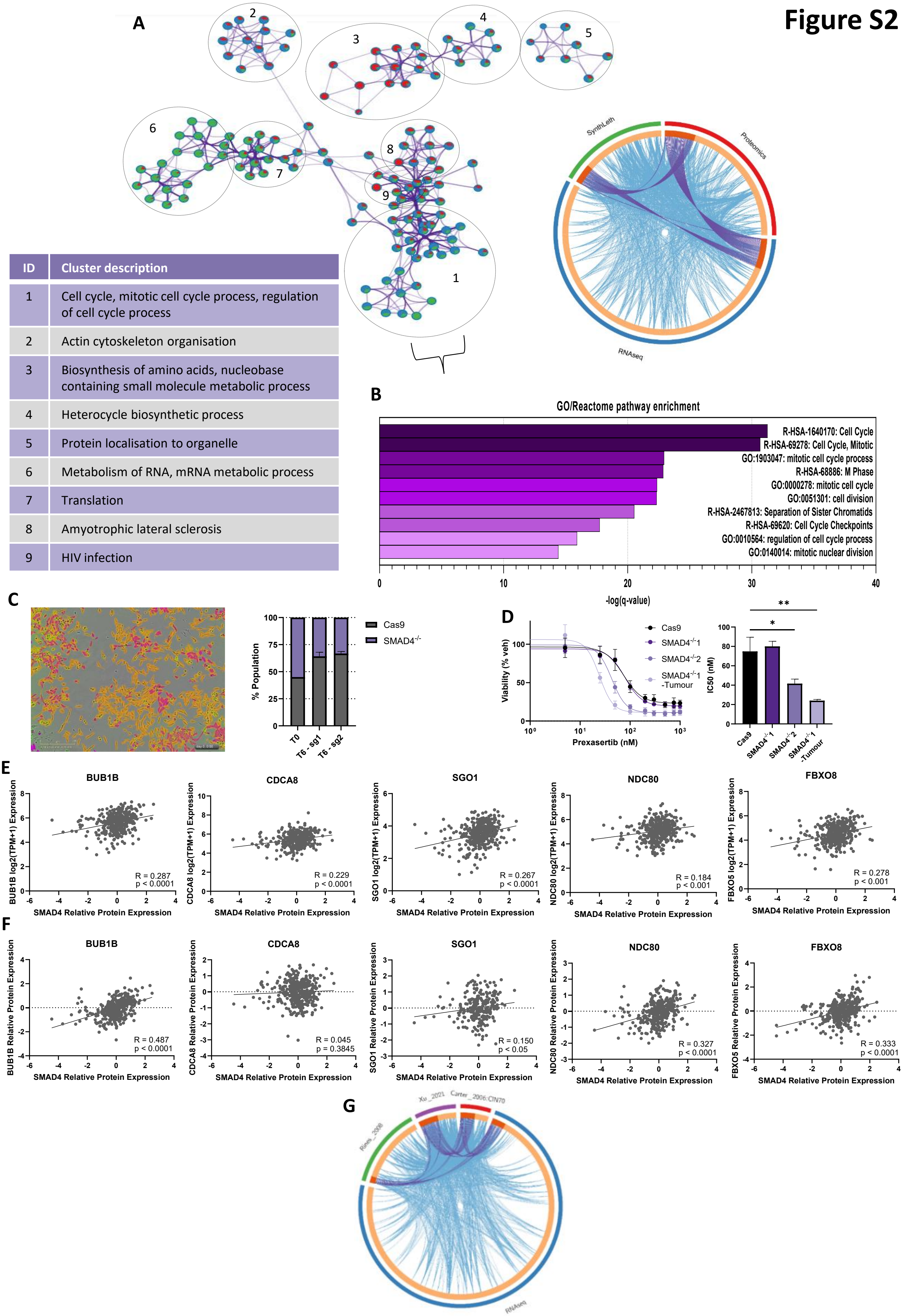


| ID | Cluster description |
| --- | --- |
| 1 | Cell cycle, mitotic cell cycle process, regulation of cell cycle process |
| 2 | Actin cytoskeleton organisation |
| 3 | Biosynthesis of amino acids, nucleobase containing small molecule metabolic process |
| 4 | Heterocycle biosynthetic process |
| 5 | Protein localisation to organelle |
| 6 | Metabolism of RNA, mRNA metabolic process |
| 7 | Translation |
| 8 | Amyotrophic lateral sclerosis |
| 9 | HIV infection |

### Figure S3

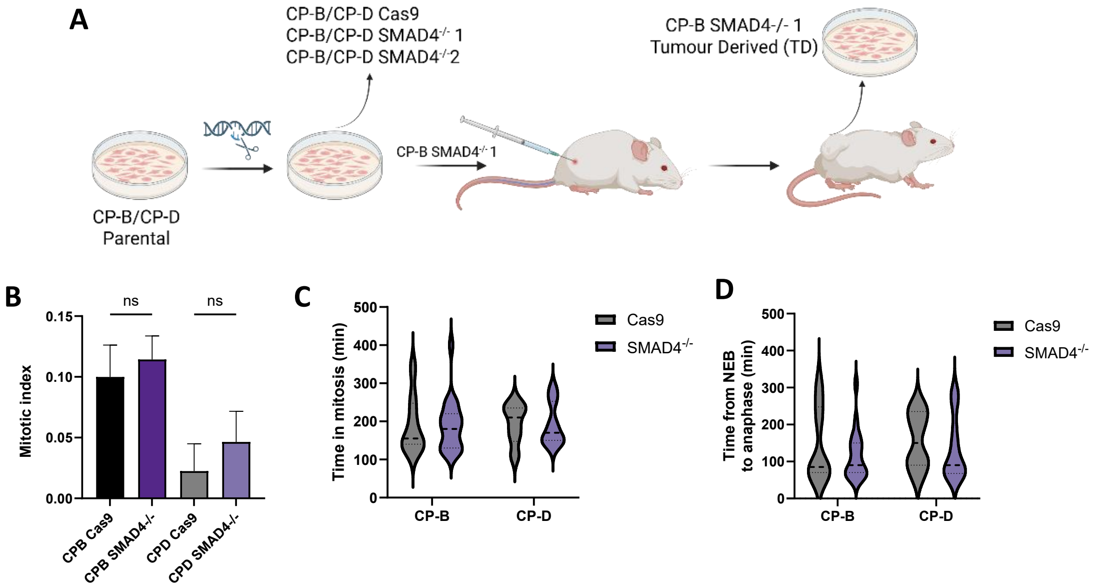
A
CP-B/CP-D Cas9
CP-B/CP-D SMAD4-/- 1
CP-B/CP-D SMAD4-/-2
CP-B SMAD4-/- 1
Tumour Derived (TD)
CP-B SMAD4-/- 1
CP-B/CP-D
Parental
B
ns
ns
Mitotic index
0.15
0.10
0.05
0.00
CPB Cas9
CPB SMAD4-/-
CPD Cas9
CPD SMAD4-/-
C
Time in mitosis (min)
500
400
300
200
100
0
CP-B
CP-D
Cas9
SMAD4-/-
D
Time from NEB
to anaphase (min)
500
400
300
200
100
0
CP-B
CP-D
Cas9
SMAD4-/-

### Figure S4

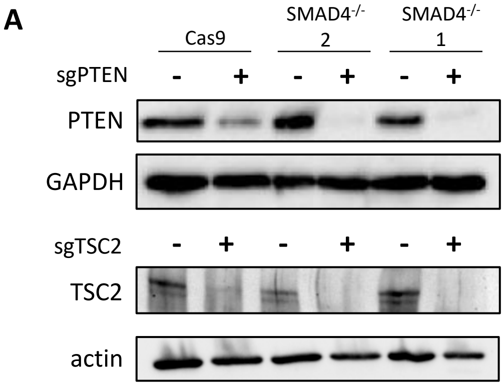
A
Cas9
SMAD4-/- 2
SMAD4-/- 1
sgPTEN
- + - + - +
PTEN
GAPDH
sgTSC2
- + - + - +
TSC2
actin

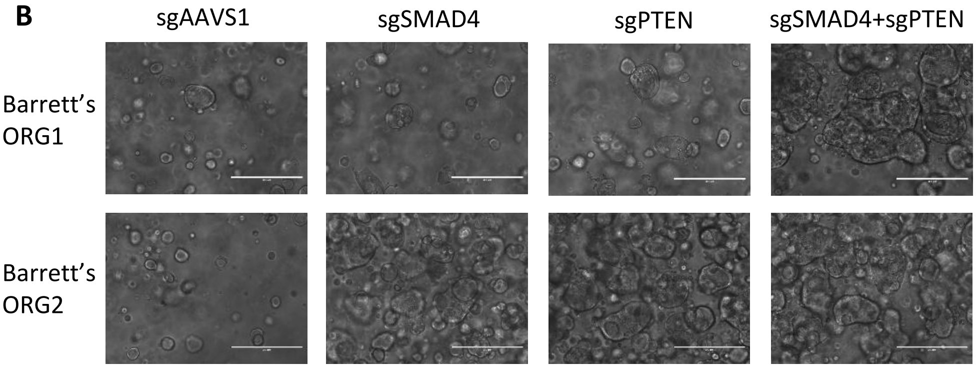
B
sgAAVS1
sgSMAD4
sgPTEN
sgSMAD4+sgPTEN
Barrett's ORG1
Barrett's ORG2

### Figure S5

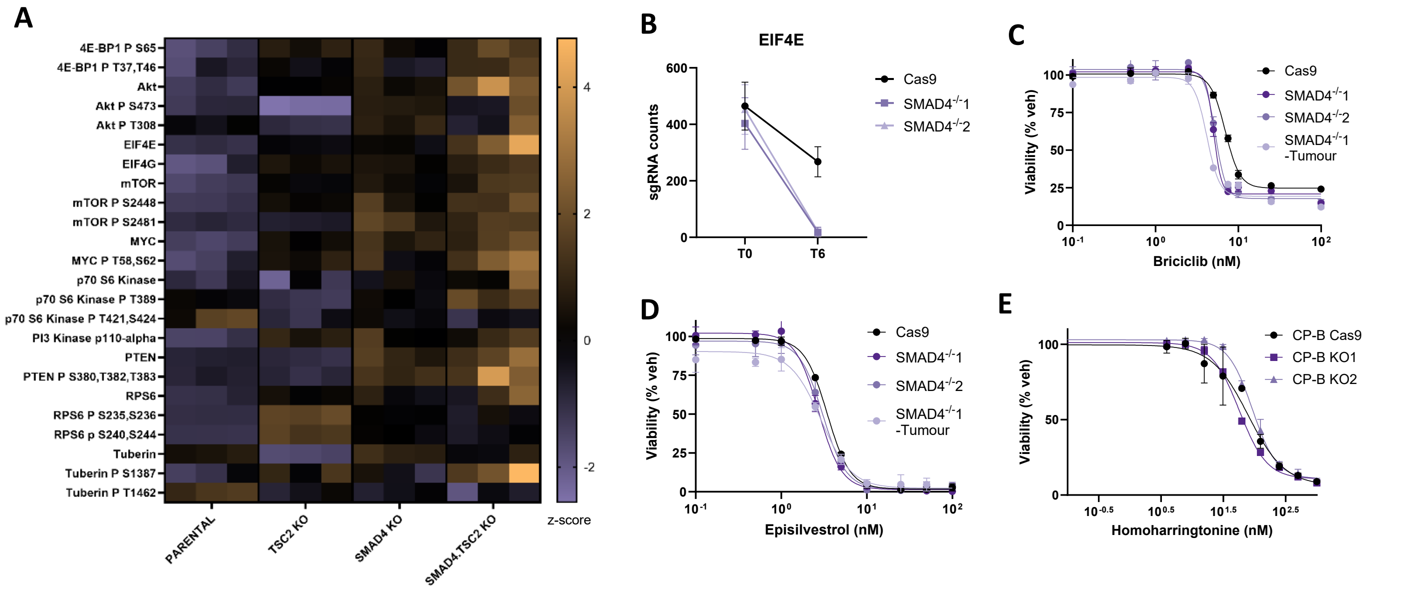

A
4E-BP1 P S65
4E-BP1 P T37,T46
Akt
Akt P S473
Akt P T308
EIF4E
EIF4G
mTOR
mTOR P S2448
mTOR P S2481
MYC
MYC P T58,S62
p70 S6 Kinase
p70 S6 Kinase P T389
p70 S6 Kinase P T421,S424
PI3 Kinase p110-alpha
PTEN
PTEN P S380,T382,T383
RPS6
RPS6 P S235,S236
RPS6 p S240,S244
Tuberin
Tuberin P S1387
Tuberin P T1462
PARENTAL
TSC2 KO
SMAD4 KO
SMAD4.TSC2 KO
z-score
B
EIF4E
sgRNA counts
T0
T6
Cas9
SMAD4-/-1
SMAD4-/-2
C
Viability (% veh)
Briciclib (nM)
Cas9
SMAD4-/-1
SMAD4-/-2
SMAD4-/-1 -Tumour
D
Viability (% veh)
Episilvestrol (nM)
Cas9
SMAD4-/-1
SMAD4-/-2
SMAD4-/-1 -Tumour
E
Viability (% veh)
Homoharringtonine (nM)
CP-B Cas9
CP-B KO1
CP-B KO2
