## Supplementary Tables S1 S2 S6 S7 for "SMAD4 loss drives chromosomal instability during tumourigenesis via translational reprogramming"

**Supplementary Table S1. Status of *MDM2* in patient samples with *SMAD4* alteration and no *TP53* alteration**

| **Study** | **Sample ID** | **SMAD4: CNA** | **SMAD4: Mutations** | **TP53: CNA** | **TP53: Mutations** | **TP53: Protein expression z-scores** | **MDM2: CNA** |
| --- | --- | --- | --- | --- | --- | --- | --- |
| **TCGA** |  |  |  |  |  |  |  |
|  | TCGA-2H-A9GI-01 | Shallow Deletion | - | Diploid | - | -0.7214 | - |
|  | TCGA-L5-A43M-01 | Shallow Deletion | - | Diploid | - | -0.7044 | Amplification |
|  | TCGA-L5-A8NF-01 | Shallow Deletion | - | Diploid | - | -0.1129 | Gain |
| **MSK** |  |  |  |  |  |  |  |
|  | P-0001323-T01-IM3 | Deep Deletion | - | Diploid | - | NA | - |
|  | P-0001542-T01-IM3 | Diploid | L540del | Diploid | - | NA | Amplification |
| **UQ** |  |  |  |  |  |  |  |
|  | SOG091_TM | Shallow deletion | G352V | Diploid | - | NA | Amplification |

**Supplementary Table S2. Criteria for designation of SMAD4 status in OAC patient samples**

| **Stratification criteria (SMAD4)** | **“Deficient” n=54** | **“Intact” n=116** | **“Strict deficient” n=28** | **“Strict intact” n=28** |
| --- | --- | --- | --- | --- |
| Log2CN (Copy Number; ASCAT) | -2 (Deletion) or -1 to -2 (LOH) | -0.5 to 0.5 (Normal) | -2 (Deletion) or -1 to -2 (LOH) | -0.5 to 0.5 (Normal) |
| Mutation | Any exonic | Absent or intronic only | Known deleterious in LOH only | Absent or intronic only |
| RNA expression z-score | < -1 | > -1 | < -1.5 | -0.5 to 0.5 |

**Supplementary Table S3 – RNAseq (differentially expressed genes)**

**Supplementary Table S4 – Proteomics (differentially abundant proteins)**

**Supplementary Table S5 – CRISPR screen**

**Supplementary Table S6. Antibodies used for immunoblotting**

| Antigen | Origin | Supplier | Item |
| --- | --- | --- | --- |
| 4EBP1 | Rabbit | CST | 9644 |
| 4EBP1 P T37/46 | Rabbit | CST | 2855 |
| CDK11 | Rabbit | CST | 5524 |
| EIF4E | Rabbit | CST | 2067 |
| EIF4G | Rabbit | CST | 2498 |
| PTEN | Rabbit | CST | 9559 |
| Puromycin | Mouse | Merck | MABE343 |
| SMAD4 | Rabbit | CST | 38454 |
| TSC2 | Rabbit | CST | 3612 |

**Supplementary Table S7. Antibodies used for RPPA**

| **Antigen** | **Origin** | **Supplier** | **Item** |
| --- | --- | --- | --- |
| 4E-BP1 P S65 | rabbit | CST | 9451 |
| 4E-BP1 P T37,T46 | rabbit | CST | 2855 |
| Akt | rabbit | CST | 9272 |
| Akt P S473 | rabbit | CST | 4060 |
| Akt P T308 | rabbit | CST | 2965 |
| EIF4E | Rabbit | CST | 2067 |
| EIF4G | Rabbit | CST | 2498 |
| mTOR | rabbit | CST | 2972 |
| mTOR P S2448 | rabbit | CST | 2971 |
| mTOR P S2481 | rabbit | Millipore | 09-343SP |
| MYC | rabbit | CST | 5608 |
| MYC P Th58,S62 | rabbit | Epitomics | 1203-1 |
| P70 S6 Kinase | rabbit | CST | 9202 |
| p70 S6 Kinase P Th89 | rabbit | CST | 9205 |
| p70 S6 Kinase P T421,S424 | rabbit | CST | 9204 |
| PI3 Kinase p110-alpha | rabbit | CST | 4249 |
| PTEN | rabbit | CST | 9552 |
| PTEN P S380,T382,T383 | rabbit | CST | 9554 |
| S6 Ribosomal Protein | rabbit | CST | 2217 |
| S6 Ribosomal protein P S235,S236 | rabbit | CST | 2211 |
| S6 Ribosomal protein P S240,S244 | rabbit | CST | 2215 |
| Tsc-2 (Tuberin) | rabbit | CST | 3612 |
| Tsc-2 (Tuberin) P S1387 | rabbit | CST | 5584 |
| Tsc-2 (Tuberin) P T1462 | rabbit | CST | 2617 |

**Supplementary Table S8. Primers used for qPCR**

| **Gene** | **Forward (5’ – 3’)** | **Reverse (5’ – 3’)** |
| --- | --- | --- |
| 4EBP1 | TATGACCGGAAATTCCTGATGGA | CCGCTTATCTTCTGGGCTATTG |
| GAPDH | GGTGTGAACCATGAGAAG | CCACAGTTTCCCGGAG |
